# Fine-scale spatio-temporal variation in natural selection on reproductive traits across two wild bird species

**DOI:** 10.64898/2026.02.27.708424

**Authors:** Jørgen S. Søraker, Yimen G. Araya-Ajoy, Ella F. Cole, Ben C. Sheldon

## Abstract

Natural selection drives adaptive evolution and underpins eco-evolutionary processes. However, the causes of variation of selection (strength and direction), and its spatio-temporally scaling, remain poorly understood. Most selection estimates are population-wide, ignoring the variation in fine-scale selection regimes within populations. We quantify variation in selection within sub-populations of a long-term study of great tits (*Parus major*) and blue tits (*Cyanistes caeruleus*) in Wytham Woods, UK. Using mixed-effect modelling, we decomposed temporal, spatial and spatio-temporal variation in selection on laying date, clutch size and their correlational selection. Across species, we detected non-linear selection for larger clutch size and earlier laying, and negative correlational selection. Spatio-temporal effects explained most variation in selection across traits and species, suggesting that the causes of selection are controlled by interacting temporal (e.g. climate) and spatial (e.g. habitat) mechanisms. Laying date had strong temporal (among-year) variation for both species, whereas spatial components generally explained little variation. Selection estimates were highly correlated between species for all traits except for correlational selection, with higher agreement when analyzing fledgling versus recruitment numbers. Our results suggest common factors underpinning small-scale spatio-temporal variation in selection in both species and that such analyses can identify the sources causing natural selection.

## Introduction

Phenotypic selection, defined as the covariance between (relative) fitness and phenotypes, is the predominant driver underlying adaptive evolution (Svensson 2023), and has been a key parameter within ecological and evolutionary research for many decades (Kingsolver et al. 2001; Siepielski et al. 2009, 2013; Svensson 2023). Phenotypic selection can shape the mean phenotype of a population (Lande and Arnold 1983; Johnson et al. 2023), therefore influencing eco-evolutionary processes like local adaptation (Savolainen et al. 2013), population divergence (Schluter 2000) and speciation (Funk et al. 2006).

Given the importance of phenotypic selection, there has long been interest in estimating selection in natural populations (Svensson 2023). The number of phenotypic selection coefficients estimates increased rapidly once Lande (1976) and Lande and Arnold (1983) explained how regression could be used to estimate linear (and quadratic) selection gradients (Kingsolver et al. 2001; Siepielski et al. 2009, 2013; Svensson 2023). This has further allowed studies of temporal and spatial fluctuation in selection (Kingsolver et al. 2001; Siepielski et al. 2009, 2013). Such fluctuating selection can have important evolutionary consequences (Lande 2007; Chevin et al. 2015), and is suggested to slow down long-term evolutionary change (Estes and Arnold 2007) due to variation in the direction and strength of selection (Bonnet and Postma 2018). While early studies considered mostly fluctuations in the direction of selection, recent studies have also acknowledged the importance of variation in the strength of selection to understand long-term evolutionary dynamics (Lande 2007; Sæther et al. 2016). Several mechanisms can drive fluctuating selection, such as changes in the population mean phenotype, changes in the optimum phenotype through environmental change, and demographic or environmental stochasticity (Engen and Sæther 2014; Chevin et al. 2015; de Villemereuil et al. 2020). Pinpointing the specific causes of variation in selection is difficult (MacColl 2011; Chevin et al. 2015; Svensson 2023), although grouping the causes into sources (e.g. demographic mechanisms, climatic patterns, spatio-temporal patterns etc.) can provide valuable information (Araya-Ajoy et al. 2025).

Although selection estimates from wild populations have generated considerable insight, they are typically estimated at population-wide scales (e.g. Sheldon et al. 2003). This implicitly assumes a homogeneous selection regime within a population, ignoring potential variation in selection due to biotic and abiotic factors that vary at different scales (Svensson and Sinervo 2004; MacColl 2011; Marrot et al. 2022). Variation in fitness, and hence selection, is caused by the environmental ecology surrounding individuals, which is likely to be constantly varying (MacColl 2011), also within populations. Hence, population-wide studies on selection are likely to yield an over-simplified view of a key element of eco-evolutionary dynamics (Govaert et al. 2019). Quantifying variation in selection at finer scales can provide better understanding of several processes. First, it can potentially inform us about the ecological drivers of selection (MacColl 2011). Second, it can improve our predictions of phenotypic change both in the short-term and in the long-term (e.g. by impacting the standing genetic variation in the population and local adaptation (Lande 2007; Quinn et al. 2009; Porlier et al. 2012). However, estimating fine-scale selection coefficients is challenging because it requires large sample sizes and defined subpopulations within which selection can be estimated and examples are scarce (see Svensson and Sinervo 2004; Lundblad and Conway 2020; Marrot et al. 2022 for exceptions). Marrot et al. (2022) explored spatial variation in selection in the common snapdragon (*Antirrhinum majus)* and showed that the strength and direction of selection can vary at scales as small as one meter (Marrot et al. 2022). Variation in selection pressures was also found to vary with elevation in a population of yellow-eyed juncos (*Junco phaeonotus*) (Lundblad and Conway 2020). Further, Svensson and Sinervo (2004) used a metapopulation of lizards to show that fine-scale local selection gradients tend to follow the overall population estimate, but substantial variation in selection regimes existed within the metapopulation.

Selection could vary both spatially (consistent differences in selection among areas), temporally (e.g. differences in selection among years for the same areas) or as a combination of the two (interacting spatial and temporal differences; Figure 1). However, very few studies decompose all three sources of variation (Araya-Ajoy et al. 2025), and most studies focus on either temporal or spatial variation in selection. Compilations of selection estimates across studies have shown that selection generally varies more spatially than temporally (Morrissey and Hadfield 2012; Siepielski et al. 2013). This is supported by quantification of within-versus among-population variation in selection gradients in exploration behaviour in great tits (Mouchet et al. 2021): the among-population variation in selection exceeded within-population estimates, but significant variation existed also within populations (Garant et al. 2007; Quinn et al. 2009; Mouchet et al. 2021). However, in general, we know little about how variation in selection within a population decomposes into spatio-temporal patterns.

**Figure 1:**
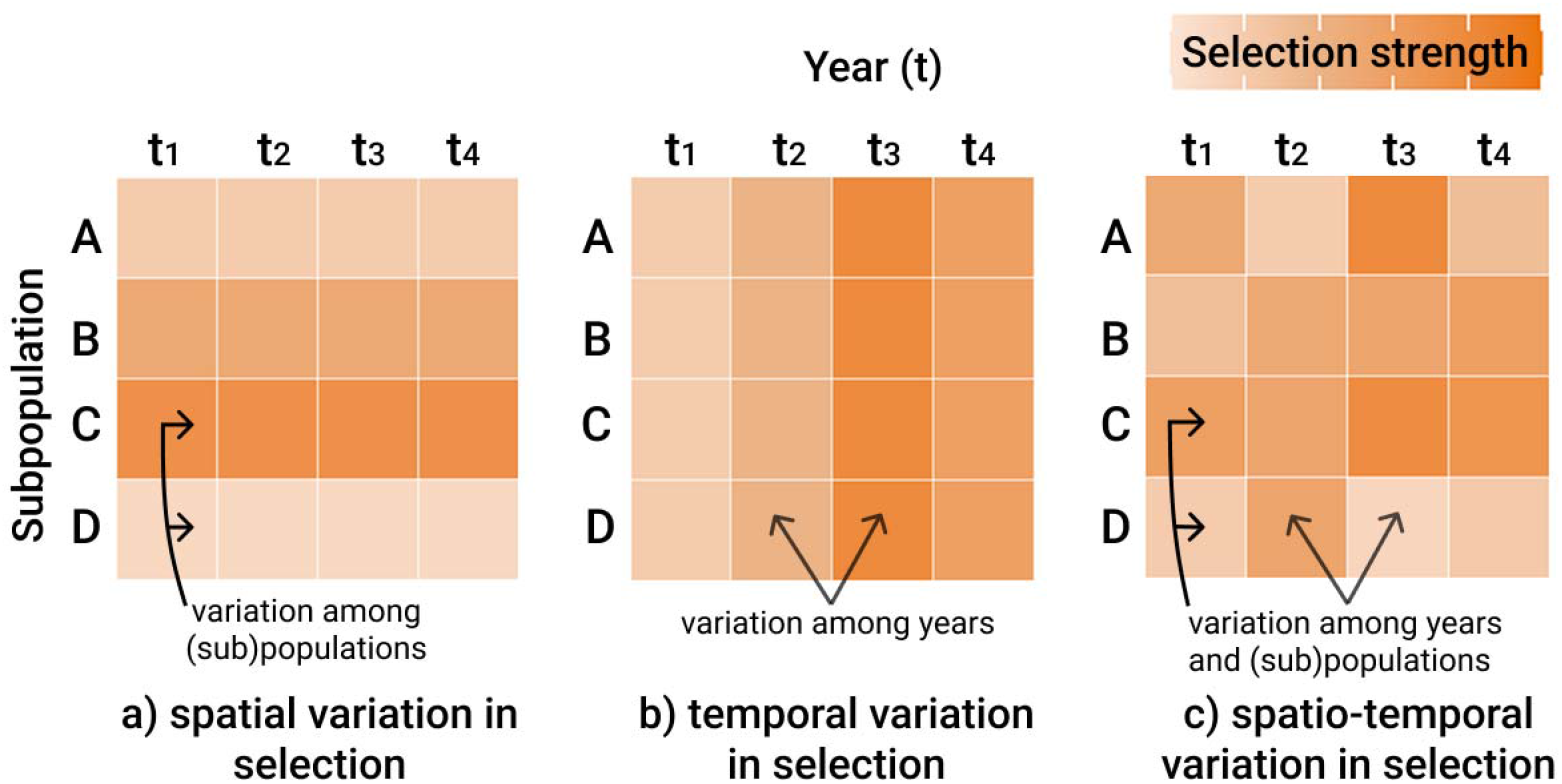
Visualization of how selection can vary a) spatially; where selection varies among subpopulations only, b) temporally; where selection is the same in all subpopulations, but are free to vary consistently among years and c) spatio-temporally; where selection varies among subpopulations and among years simultaneously.

The relationship between fitness and reproductive traits, like the time of breeding onset and clutch size in birds, has been frequently addressed by field studies of natural selection (Garant et al. 2007). Breeding onset and clutch size are key reproductive traits (Perrins 1965; Perrins and McCleery 1989; Shave et al. 2019), and studies on selection of these traits are plentiful. Such studies often find selection for earlier breeding onset (Garant et al. 2007; Radchuk et al. 2019; Shave et al. 2019; Branston et al. 2021; Jantzen and Visser 2023), which can be caused by variety of different ecological drivers, such as seasonal variation in food availability (Perrins and McCleery 1989; Visser and Gienapp 2019), predation rates (Perrins 1965) and post-fledging competitive advantage for early-fledged young (Nilsson and Smith 1988). These processes may also vary at fine scales. For example, laying date has been shown to depend on the tree phenology close to the nestbox in a population of great tits (Hinks et al. 2015), with tree phenology being highly heterogeneous, but individually consistent, across the population (Hinks et al. 2015; Cole and Sheldon 2017). Selection for larger clutch size is also documented (Garant et al. 2007; Branston et al. 2021). However, although laying date and clutch size show strong phenotypic correlation (Suárez et al. 2005; Garant et al. 2007; Vaugoyeau et al. 2016; Vriend et al. 2023), few studies have estimated their correlational selection (i.e. selection on the combination of traits; (Sheldon et al. 2003; Garant et al. 2007)). Most studies on these traits, however, investigate selection on a population-based level (Visser and Holleman 2001; Sheldon et al. 2003), again ignoring potential fine-scaled selection regimes.

This study had three objectives. First, we aimed to study how selection on two reproductive traits, laying date and clutch size, as well as their correlational selection, varied within populations of great tits and blue tits in Wytham Woods, UK. This also allowed us to decompose the variation in selection regimes into temporal, spatial and spatio-temporal components. A previous study from this population demonstrated that differences in selection regimes contrasted between east and west sections of this population (Garant et al. 2007), suggesting that there could be small-scale processes affecting selection. Second, we aimed to explore how edge-effects influence selection regimes for both traits. Previous investigations from this population show that birds near edges produce fewer recruits, express later laying dates and produce smaller clutches with larger eggs (Wilkin et al. 2007). However, whether the selection regimes vary between edge-territories and central territories and therefore contributes to spatial variation in selection remains unknown. Lastly, we aimed to investigate the correlation in selection estimates across these two species. While they are often assumed to be similar in terms of their ecology, tests of whether they experience similar selection regimes at given time and location is lacking, despite numerous studies on both species (Garant et al. 2007; Porlier et al. 2012). The similarity in selection across species for a given time and location yields insight into the extent that selection varies due to species-specific versus other effects.

## Methods

### Data collection

The data were collected in Wytham Woods, near Oxford, UK (51°46′ N, 1°20′ W). The woods consist of mixed-deciduous woodland with surrounding farmland (Savill et al. 2010). The great tit and blue tit populations have been studied here since 1947, mainly through breeding events in nest boxes (Perrins 1965; Harvey et al. 1979). Nests are monitored throughout the breeding season (at least once per week). Date for first egg laid, clutch size, and number of fledglings are recorded for all broods. The nestlings are ringed at the age of 14 days with a unique metal ring from the British Trust of Ornithology. This enables us to monitor recruitment of fledged individuals to the breeding population and their following reproductive traits. Adults are further caught and identified at the nest during breeding.

### Statistical analysis

We use the number of fledglings and recruits (fledglings that survive and join the breeding population in subsequent years, and breed in a monitored nestbox in the study population) as measures of fitness. The first egg date was used as laying date and the maximum egg number as clutch size. For clutch size, we included only clutches where at least one egg hatched to exclude uncompleted clutches that failed during laying. We restricted the analysis to only include first clutches, determined as the first brood of the identified female each year. Hence, first clutches could include both replacement clutches and second clutches by unidentified females (e.g. breeding failed before they could be identified). However, true second clutches are rare in this population, and normally appear much later in the season (Perrins and McCleery 1989). We also excluded clutches subject to experimental manipulations, since they affected both measures of fitness. The study period was restricted to the years 1980 and onwards, as this marked a change in the type of nest boxes used due to heavy predation in previous years; previous variable predation rates are known to have impacted the phenotypic distribution of clutch size in particular (e.g. Julliard et al. 1997). The last year in which data were analysed was 2023 (with recruitment to 2024). For blue tits, the years 1991-2000 were excluded due to limited fledgling ringing in this period. The study thus spanned 44 years and 10477 breeding attempts for great tits while spanning 34 years and 8986 breeding attempts for blue tits. Statistical analysis was performed in R, version 4.3.1 (R Core Team 2018), mainly by using the “lme4” (Bates et al. 2014) and “rstan” (Stan Development Team 2016) packages. When models suffered from overdispersion, we corrected for this by using a negative binomial model in the “glmmTMB” package (Brooks et al. 2017).

### Sub-populations

To investigate spatial, temporal and spatio-temporal variation in selection, one needs long-term data for several defined restricted areas (populations). To do this, we split the population into 8 subpopulations (Figure S1, number of breeding events per subpopulation-year given in Table S1 & S2; subpopulation P was excluded from the analysis due to consistently low sample size). Although these subpopulations clearly form part of a single metapopulation, they exhibit several consistent differences across the course of the period data collection. First, there is up to a five-fold difference in density of nest-boxes, and a previous study highlighted that these subpopulations show variation in population size growth rates similar to that among populations (Gamelon et al. 2019). Moreover, they vary in habitat types (based on habitat analysis in previous studies; Gibson 1988), altitude (Table S3), and they can also vary in population density (measured as the inverse of territory size) (Table S4 & S5). Besides the differences in nest-box density, we had no *a priori* reason to expect different selection regimes, though we know they differ in environmental composition, a feature we argue is common within most study systems. However, there was relatively little variation in laying date and clutch size among the subpopulations (Figure S2-5), and the subpopulation described little of the total variance in both traits for both species (Table S6). Both laying date and clutch size was standardized so that each trait was measured in number of standard deviations (SD) from the mean within each subpopulation within each year (that is, scaled to a mean of 0 and variance of 1). Since selection estimates can be sensitive to the standard deviation values, particularly when population size is small, we used the standard deviation for each subpopulation over the entire study period instead of the standard deviation for the given subpopulation for the given year. Blue tits and great tits were analysed separately.

Since sampling error can bias the estimation of variation in selection, we estimated selection gradients using a mixed effect modelling approach (Morrissey and Hadfield 2012). We used a random regression mixed effect model to estimate the linear and quadratic effects of laying date and clutch size on fitness components, as well as the interaction between the linear effects. We assumed a Poisson error distribution and used a log-link function, which effectively models the effects of clutch size and laying date on the expected log fitness. This allowed estimation of the parameters of a multivariate Gaussian fitness function (Morrissey and Goudie 2022). We model random intercepts for subpopulation, year and subpopulation-year (combination of subpopulation and year). We also fitted random slopes for the linear fixed effects of the traits on fitness; for mathematical notation, see Supplementary material, equation 1. By including random slopes, we can decompose the variation in selection associated to spatial, temporal and spatio-temporal heterogeneity (calculation description in figure S6). The estimates for the fixed effects represent the overall selection estimate across all the subpopulations across all years. Assuming that all subpopulations experience the same strength of stabilizing selection (equal values of the non-linear components), we also estimated the optimum phenotype for each subpopulation for each year by dividing the linear selection estimate by two times the negative quadratic selection estimate. We thereafter calculated the variances on mean phenotypes and optimums for both subpopulation and population level estimates over time.

To check if the proportion of variation in selection was driven by differences in mean phenotypes among the subpopulations rather than changes in the optimum, we also performed the same model as above, but standardizing each trait to the population mean rather than the subpopulation mean. This reduces the variation in mean phenotype both spatially and temporally. We also checked if female identity could affect our results by including this as a random effect in an additional model otherwise structured as above. Initially, all models were run without female identity, as females with earlier laying dates are less likely to be detected (Kidd et al. 2015). When using the number of recruits as fitness measurement, selection estimates could be biased if the trait of interest co-varies with the number of birds that disperse out of the population. To investigate if such dispersal could affect our selection estimate, we used a small subset of known recoveries outside of our population and modelled the number of dispersers by the same fixed effects as above. In addition, we included the same random effects but did not allow for random slopes due to low sample sizes (for details about this subset of dispersal data, see supplementary materials:Dispersal). Recruitment dispersal out of the population could result in a small underestimation of selection on clutch size, quadratic clutch size and correlational selection for great tits, as these traits were statistically significantly associated with dispersal propensity in our limited dispersal dataset (Table S7). However, no such trends were found for blue tits. Including mother identity in the models provided similar results (but due to reduced sample size and bias in laying date for detected and identified females, we have placed these analyses in the supplementary materials (Table S8)). Moreover, the frequentist approach of the same model provided similar results as the Bayesian model.

Regarding variation in selection, by dividing the random slope component (variation in selection explained by a source) by the total variation in selection (all random slope components combined), we obtained the proportion of variation in selection explained by the different factors (hence, the percentages should sum to approximately 100%). In this way, we decomposed the variation in selection estimates due to spatial, temporal and spatio-temporal sources. However, obtaining proper uncertainty estimates of these proportions are challenging. Therefore, we constructed a Bayesian mixed-model using Stan through RStan (Stan Development Team 2016) with the same model structure as above. We calculated each proportion of variance explained by each of the random effects (year, subpopulation and subpopulation-year) as a generated quantity in the model. In this way, we obtain one posterior distribution for each of the proportion calculations, from which we can infer the uncertainty of these proportions. The models were run with two chains, 600 burn-ins and iterations adjusted to the conversion of the r-hat statistic (Gelman et al. 1995). This Bayesian modelling approach extracts the median of the posterior distributions, which could result in the three components not summing exactly to 100%. Hence, small deviations from 100% may occur.

### Edge-territories

To investigate whether selection differed between edge-territories and central territories, we assigned an edge-status to every occupied nest-box for each year. This was done by constructing Theissen-polygons around each occupied nestbox. In these polygons, we use the GPS-location of each occupied nest-box and determine all areas that are closest to this specific nestbox compared to the other occupied nestboxes (Wilkin et al. 2007a,b). This area was used as a measure of territory size. By identifying the outline of the forest, the nest-boxes with Theissen-polygons that include this outline for can be identified as edge-territories while those without the wood outline can be identified as central territories. Edge-territories are generally larger (i.e. lower density) (Table S9) but have similar numbers of oaks as central territories (Table S10).

When each occupied nest-box for each year was assigned a territory, we further created an edge-status-year combination (on the edge of the population for that year or not). We then standardized laying date and clutch size within each edge-status-year combination to have a mean of 0 and variance of 1. We used two linear mixed-models with Poisson error distribution and a log-link function to model (the log) number fledglings and (the log) number of recruits as response variables, respectively. We used laying date, quadratic laying date, clutch size, quadratic clutch size and edge-status as fixed effects. Additionally, we incorporated the interactions between edge-status and laying date, between edge-status and quadratic laying date, between edge-status and between clutch size and last edge-status and quadratic clutch size. Year and edge-status-year were used as random effects. We further modelled random slopes for the directional selection estimate of laying date and clutch size to within both edge-status-year and year (for mathematical notation, see Supplementary materials, equation 2). The same principle of variation decomposition among the random effects follows from the description above (for subpopulations). We also checked if female identity could affect our results by including this as a random effect. However, including mother identity reduced the sample size of the model. Again, we investigated if our selection estimates could be biased when using recruit number as fitness by estimating how the number of dispersers were affected by laying date and clutch size (and their quadratic component) in addition to edge-status.

## Results

### Spatial, temporal and spatio-temporal variation in selection

#### Great tit

For great tits, when using the number of recruits as fitness measurement (hereafter recruitment selection), the population generally showed non-linear selection for earlier laying date (the general selection for the whole population; Figure 2a & S7, Table 1 & S11). Earlier initiated clutches generally had higher fitness, but the effect is non-linear, with recruit production is maximized at 1.08 standard deviations (SD) earlier than the mean, equivalent to 7.8-9.8 days earlier than the mean, depending on the standard deviation of the given subpopulation. We also found non-linear selection for larger clutches (Figure 2b, Table 1 & S11), with maximized recruit production at 0.13 SD above the mean (equivalent to 0.2-0.3 eggs more than the mean clutch size (mean=8.5 eggs), depending on the subpopulation SD). Finally, we found negative correlational selection between clutch size and laying date, illustrating selection on a combination of these traits (Table 1 & S11).

**Table 1:**
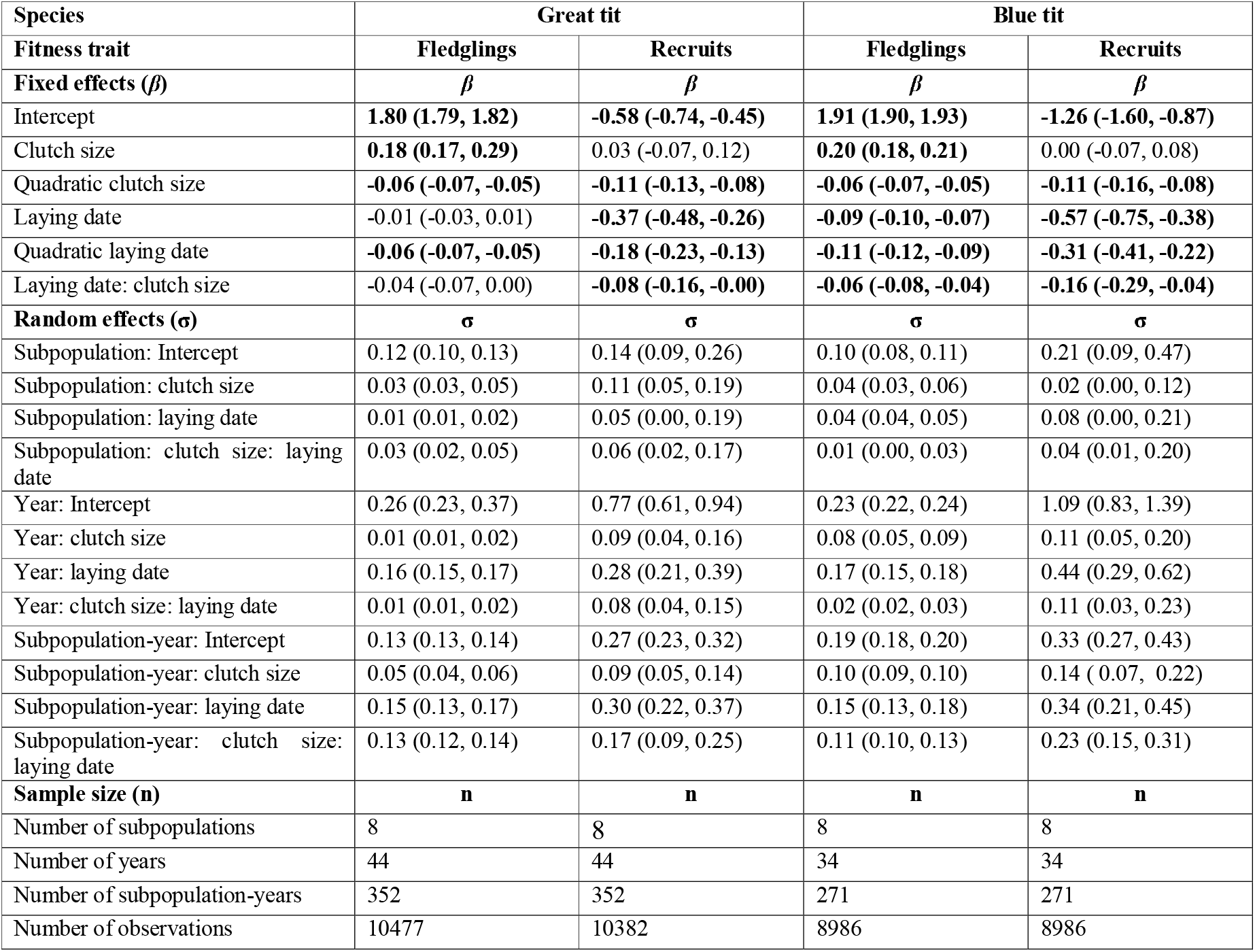
The effect of laying date, the quadratic component of laying date, clutch size, the quadratic component of clutch size, and the correlational selection of clutch size and laying date when modelling the number of fledglings and number of recruits, respectively, for both great tit and blue tit. The table is based on a Bayesian model, and the estimates represent a point estimate of the median posterior distribution and the 95% credible intervals. Followed are the estimates for the random effects, including the random slopes for clutch size and laying date within each random effect.

**Figure 2:**
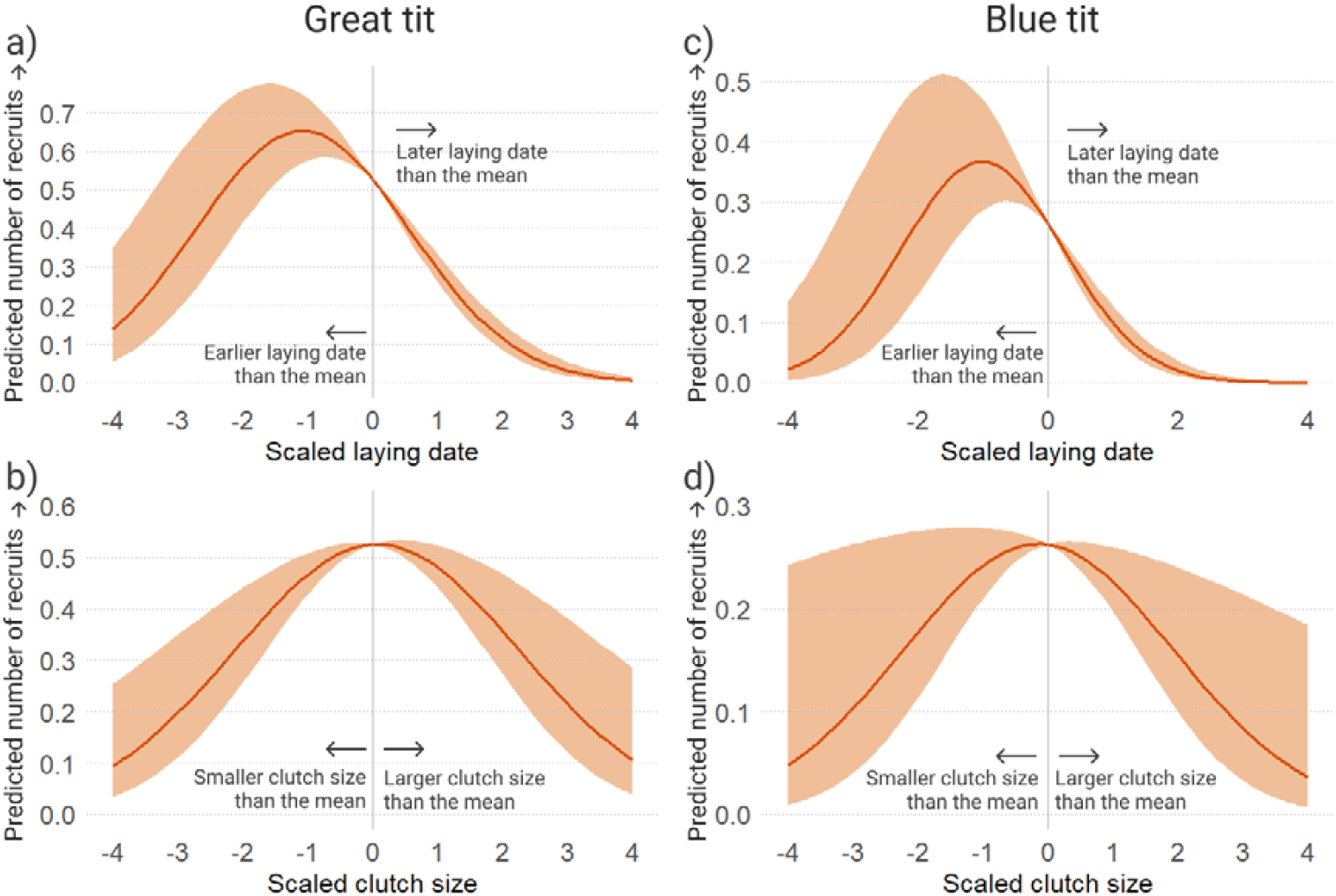
The relationship between scaled laying date and the number of recruits produced for a) great tit and b) blue tit and between scaled clutch size and the number of recruits produced for c) great tit and d) blue tits. Data are derived from long-term population study in Wytham Woods, UK, from 1980-2025. Each trait is measured in the number of standard deviations from the mean. The shaded area represents the CI of the estimated lines. Data points are excluded from this figure to facilitate interpretation (see Figure S7 and Figure S14 for data points included).

Decomposing the variance in selection to spatial, temporal and spatio-temporal components revealed that the temporal and spatio-temporal components independent-ly explained substantial parts of the variation in selection estimates on laying date (46% temporal and 50% spatio-temporal) (Figure 3a and Table 2). However, the uncertainty estimates of these parameters were wide, and the credible intervals of the spatio-temporal and the temporal components overlapped. Only a small proportion of variation was explained by the spatial subpopulation-effect (3%), suggesting little consistent variation in selection regimes among different subpopulations. For clutch size, the three sources explained similar amounts of variation (spatio-temporal (31%), temporal (29%) and the spatial component (39%; Figure 3b and Table 2). Again, the credible intervals overlapped for all three sources. For correlational selection, most variation was explained by the spatio-temporal component (71%), followed by a modest temporal year component (15%) and a small spatial component (8%) (Figure 3c and Table 2). We present the median of the posterior distributions; therefore, the proportion of the three sources does not always add to one.

**Table 2:**
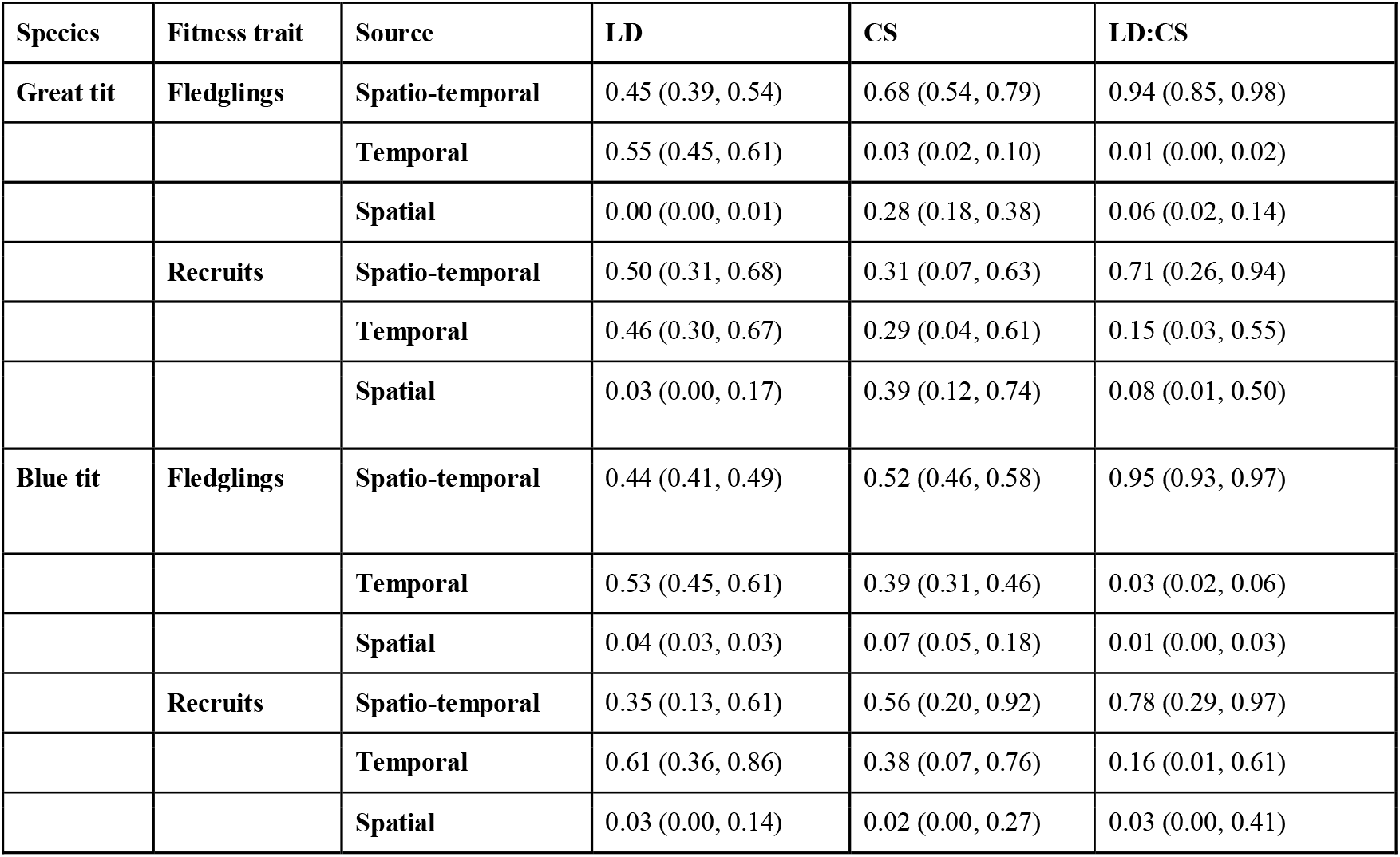
The proportion of variation in selection for the different traits (CS= clutch size, LD = laying date and LD:CS=interaction between laying date and clutch size) explained by spatio-temporal, temporal and spatial components for the two models treating fledgling number and recruit number as fitness, respectively. The parameters are based on a Bayesian model using the “Stan”-package. Presented here are the posterior medians and the 95% credible intervals. While the values for each trait per fitness trait and species should sum to 1, these results are based on a Bayesian approach and may therefore deviate slightly from this sum.

| Species | Fitness trait | Source | LD | CS | LD:CS |
| --- | --- | --- | --- | --- | --- |
| Great tit | Fledglings | Spatio-temporal | 0.45 (0.39, 0.54) | 0.68 (0.54, 0.79) | 0.94 (0.85, 0.98) |
|  |  | Temporal | 0.55 (0.45, 0.61) | 0.03 (0.02, 0.10) | 0.01 (0.00, 0.02) |
|  |  | Spatial | 0.00 (0.00, 0.01) | 0.28 (0.18, 0.38) | 0.06 (0.02, 0.14) |
|  | Recruits | Spatio-temporal | 0.50 (0.31, 0.68) | 0.31 (0.07, 0.63) | 0.71 (0.26, 0.94) |
|  |  | Temporal | 0.46 (0.30, 0.67) | 0.29 (0.04, 0.61) | 0.15 (0.03, 0.55) |
|  |  | Spatial | 0.03 (0.00, 0.17) | 0.39 (0.12, 0.74) | 0.08 (0.01, 0.50) |
| Blue tit | Fledglings | Spatio-temporal | 0.44 (0.41, 0.49) | 0.52 (0.46, 0.58) | 0.95 (0.93, 0.97) |
|  |  | Temporal | 0.53 (0.45, 0.61) | 0.39 (0.31, 0.46) | 0.03 (0.02, 0.06) |
|  |  | Spatial | 0.04 (0.03, 0.03) | 0.07 (0.05, 0.18) | 0.01 (0.00, 0.03) |
|  | Recruits | Spatio-temporal | 0.35 (0.13, 0.61) | 0.56 (0.20, 0.92) | 0.78 (0.29, 0.97) |
|  |  | Temporal | 0.61 (0.36, 0.86) | 0.38 (0.07, 0.76) | 0.16 (0.01, 0.61) |
|  |  | Spatial | 0.03 (0.00, 0.14) | 0.02 (0.00, 0.27) | 0.03 (0.00, 0.41) |

**Figure 3:**
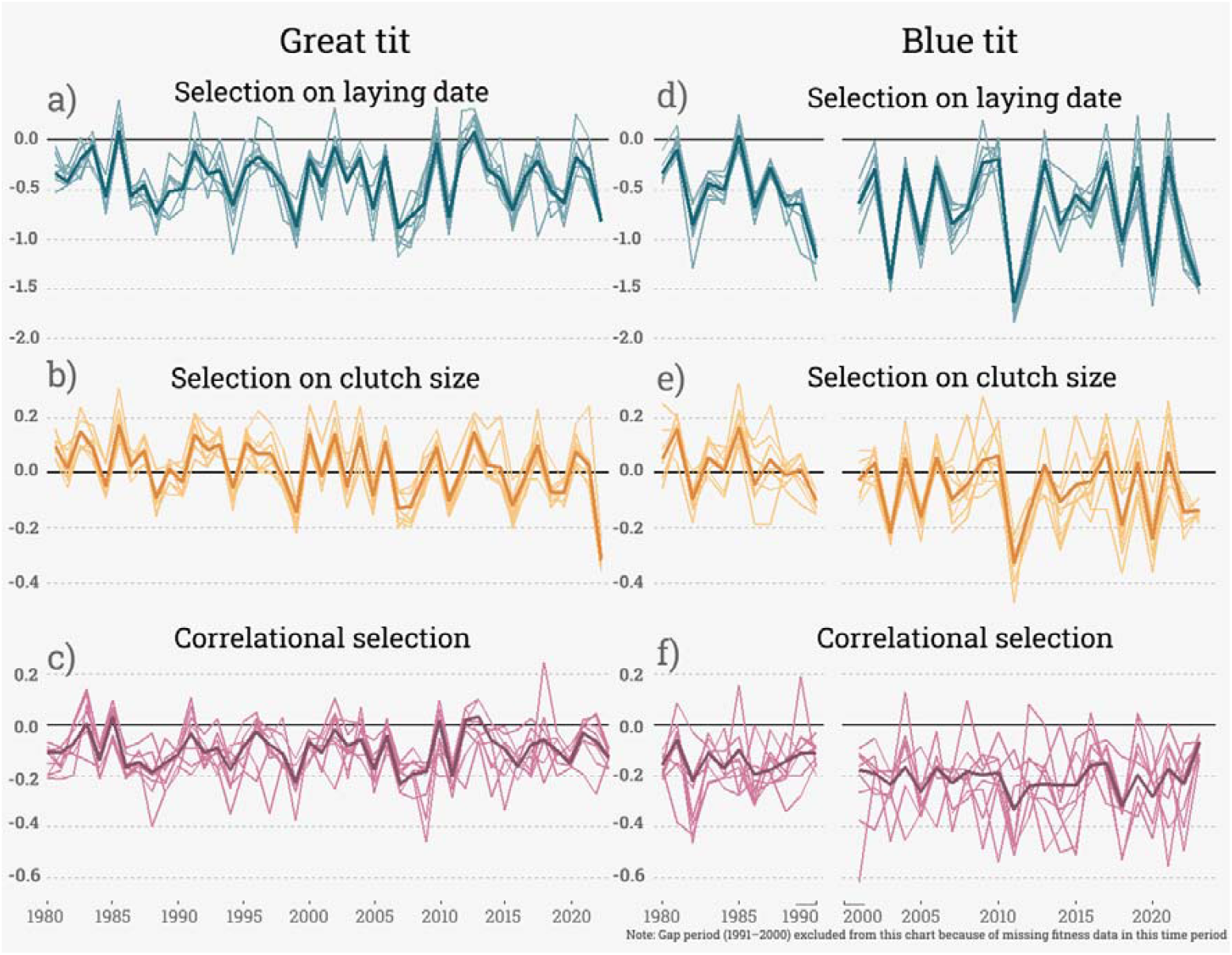
Yearly overview of the selection estimates for a) laying date in great tit, b) clutch size in great tit, c) correlational selection in great tit, d) laying date in blue tits, e) clutch size in blue tit and f) correlational selection in blue tit, when using the number of recruits as fitness measurement. Data are derived from long-term population study in Wytham Woods, UK, from 1980-2025. Each line with the weaker color represents a subpopulation while the line with stronger color represents the population-wide selection estimate. To extract the selection estimate for a given subpopulation for a given year, we combined the model estimate for the trait of interest with the estimate for that trait’s random slope of year, subpopulation and subpopulation-year. By adding these together, one obtains the overall selection estimate for each subpopulation for each year. The population-wide estimate was extracted by combining the model estimate for the trait of interest with the estimate for that trait’s random slope of year. Each trait has its own y-scale based on the variation in selection for each trait.

When considering the absolute variation in selection gradients, both laying date and clutch size selection frequently changed direction, bur more often for clutch size than laying date (Figure 3a, 3b and S8), while it remained stable for correlational selection (negative selection). Similar results were produced when estimating (variation in) selection by standardizing the phenotypes by the “whole” population mean for each year (Table S12), suggesting a limited effect of differences in mean phenotypes among the subpopulations. This is also supported by the similar distribution of both laying date and clutch size among the subpopulations (Figures S2-5). Since the subpopulation-years was assumed to have the same non-linear (quadratic component), variation in estimated optima was driven by variation in the linear selection estimates (Figure S9). There was considerable variation both in the mean phenotypes and the optimum phenotypes across the study period (Figure S9, Table S13). Notably, there appeared to be more variation in the optima than the mean phenotypes (Table S13).

When we used the number of fledglings as a fitness measurement (hereafter fledgling selection), we found little evidence for overall selection for earlier laying date, but rather a quadratic-component of laying date (Table 1; Figure S10), indicating lower fitness for earlier and later clutches compared to clutches closer to the mean laying date, with fledgling number maximized at 0.25 SD (2.0-2.3 days depending on the subpopulation SD) earlier than the mean. We also identified non-linear selection for larger clutch sizes, with the number of fledglings maximised at 1.58 SD (2.9-3.2 eggs depending on sub-population SD) above the mean (mean=8.5 eggs; Table 1; Figure S11 & S12). Hence, we found strong selection for larger clutch size when looking at fledgling production, but not for recruit production. Again, our results support a negative interaction between clutch size and laying date, representing a negative correlational selection between these two traits (Table 1). We found the correlational selection to be weaker than when we used the number of recruits as fitness measurement.

For laying date, 45% of the variation in the linear selection estimate was explained by the spatio-temporal component while 55% was explained by the temporal component (Table 1). Again, the uncertainty estimates were overlapping for these components. Only a small proportion (less than 1%) of the variation in selection estimates was explained by consistent variation among the different subpopulations. For clutch size, most variation was explained by the spatio-temporal interaction (68%) compared to relatively small temporal (3%) and moderate spatial (28%) sources. The uncertainty estimate of the two latter sources did not overlap with that of the spatio-temporal source, indicating selection to vary mostly due to spatio-temporal interactions. For correlational selection, most of the variation in selection was decomposed into the spatio-temporal factor (94%), followed by a small spatial (6%) and temporal (1%) components (Table 1).

#### Blue tit

For blue tits, when considering recruit selection, we found strong non-linear selection for earlier laying, with fitness maximised at 1.02 SD earlier than the mean (being 8.3-9.6 days earlier than the mean depending on the SD values for the subpopulations; Figure 2c, 3d & S13 and Table 1 & S11). We also found a non-linear (quadratic) component of selection on clutch size, but not a linear selection component (maximum fitness reached at mean clutch size; Figure 2d, 3e & S14). As for great tits, we found correlational selection between laying date and clutch size (Figure 3f and Table 1 & S11).

Decomposing variation in recruitment selection estimates showed that most variation in laying date selection was due to the temporal-(61%), and the spatio-temporal component (35%) (Table 2), indicating that most variation in selection was due to different conditions among the years and to interactions between sub-populations and years. The uncertainty estimates of those parameters are overlapping. Only a small proportion (3%) was due to spatial differences (Table 2). As for great tits, the absolute selection gradients varied between positive and negative at a high rate in both clutch size and laying date selection, while it was more stable negative for correlational selection (Figure S15). For clutch size selection and correlational selection in blue tits, most variation was decomposed into a spatio-temporal component (56% and 78%, respectively; Table 2). For clutch size, the temporal component explained a considerable proportion of variation in selection (38%) (Table 2), while this percent was moderate for correlational selection (16%). Only a small proportion of variation was due to consistent differences among the sub-populations (2% for clutch size and 3% for correlational selection).

When estimating fledgling selection in blue tits, we found non-linear selection for earlier laying dates, with fitness maximised at 0.54 SD earlier than the mean (4.4-5.1 days depending on the subpopulation SD). Also, we show that there is negative correlational selection on laying date and clutch size. We also show that clutch size is under non-linear selection for larger clutches (fledgling production maximised at 1.6 SD (3.0-3.3 eggs depending on the subpopulation SD) above the mean (mean=9.8 eggs), Table 1 & Figure S16-S18). As with for great tits, the correlational selection gradient was smaller when modelling fledgling numbers compared to modelling recruit numbers (Table 1 & S11).

The decomposition of variation in fledgling selection estimates showed that the spatio-temporal year component explained considerable variation for all traits (44% for laying date, 52% for clutch size and 95% for correlational selection). This was followed by the temporal interaction for all traits (53% for laying date, 39% for clutch size and 3% for correlational selection). Further, the spatial component accounted for close to no variation in selection (4% for laying date, 7% for both clutch size and 1% for correlational selection, Table 2).

#### Comparing selection estimates between species

When comparing the selection estimates for laying date, clutch size and correlational selection between the two species for each subpopulation-year combination, we found that both laying date and clutch size were moderately positively correlated when using recruit number as the fitness measure (correlation ranging from 0.303 to 0.479 for laying date and 0.073 to 0.521 for clutch size (Figure 4, Table 3)). Hence, stronger selection estimates for one species are associated with higher selection estimates in the other species, suggesting that these two species are under moderately similar (ecologically driven) selection regimes, even at this relatively fine scale. However, for correlational selection, the agreement between the selection estimates was generally lower (ranging from -0.212 to 0.327 (Table 3)). Hence, selection on the combination of laying date and clutch size seems to be much less consistent between the species than the selection for each trait alone. When using fledgling numbers as fitness, we found an even stronger agreement in estimates of laying date and clutch size selection estimates but again, little agreement in estimates of correlational selection (Table 3).

**Table 3:** The Pearson correlation estimate (*ρ*) in selection estimates between great tit and blue tit in the different subpopulations of Wytham for the different traits (CS= clutch size, LD = laying date and LD:CS=interaction between laying date and clutch size) when using fledgling number and recruit numbers as fitness. The selection estimates are extracted from a frequentist modelling approach (using the “lme4”-package). For both species, each year in both the general population and all subpopulations has a selection estimate. The correlation is based on these selection estimates.

| **Fitness trait** | **Fledglings** |  |  | **Recruits** |  |  |
| --- | --- | --- | --- | --- | --- | --- |
| **Plot** | **$\rho$ LD** | **$\rho$ CS** | **$\rho$ CS:LD** | **$\rho$ LD** | **$\rho$ CS** | **$\rho$ CS:LD** |
| Population | 0.739 | 0.741 | -0.083 | 0.495 | 0.463 | 0.099 |
| b | 0.723 | 0.711 | -0.226 | 0.479 | 0.495 | -0.191 |
| c | 0.711 | 0.642 | 0.293 | 0.357 | 0.073 | -0.007 |
| cp | 0.644 | 0.684 | -0.103 | 0.444 | 0.483 | -0.212 |
| ex | 0.618 | 0.679 | -0.077 | 0.303 | 0.430 | 0.327 |
| mp | 0.721 | 0.689 | -0.019 | 0.457 | 0.435 | 0.274 |
| o | 0.669 | 0.591 | 0.075 | 0.474 | 0.381 | 0.173 |
| sw | 0.824 | 0.728 | 0.216 | 0.377 | 0.521 | -0.105 |
| w | 0.654 | 0.516 | -0.003 | 0.356 | 0.261 | 0.207 |

**Figure 4:**
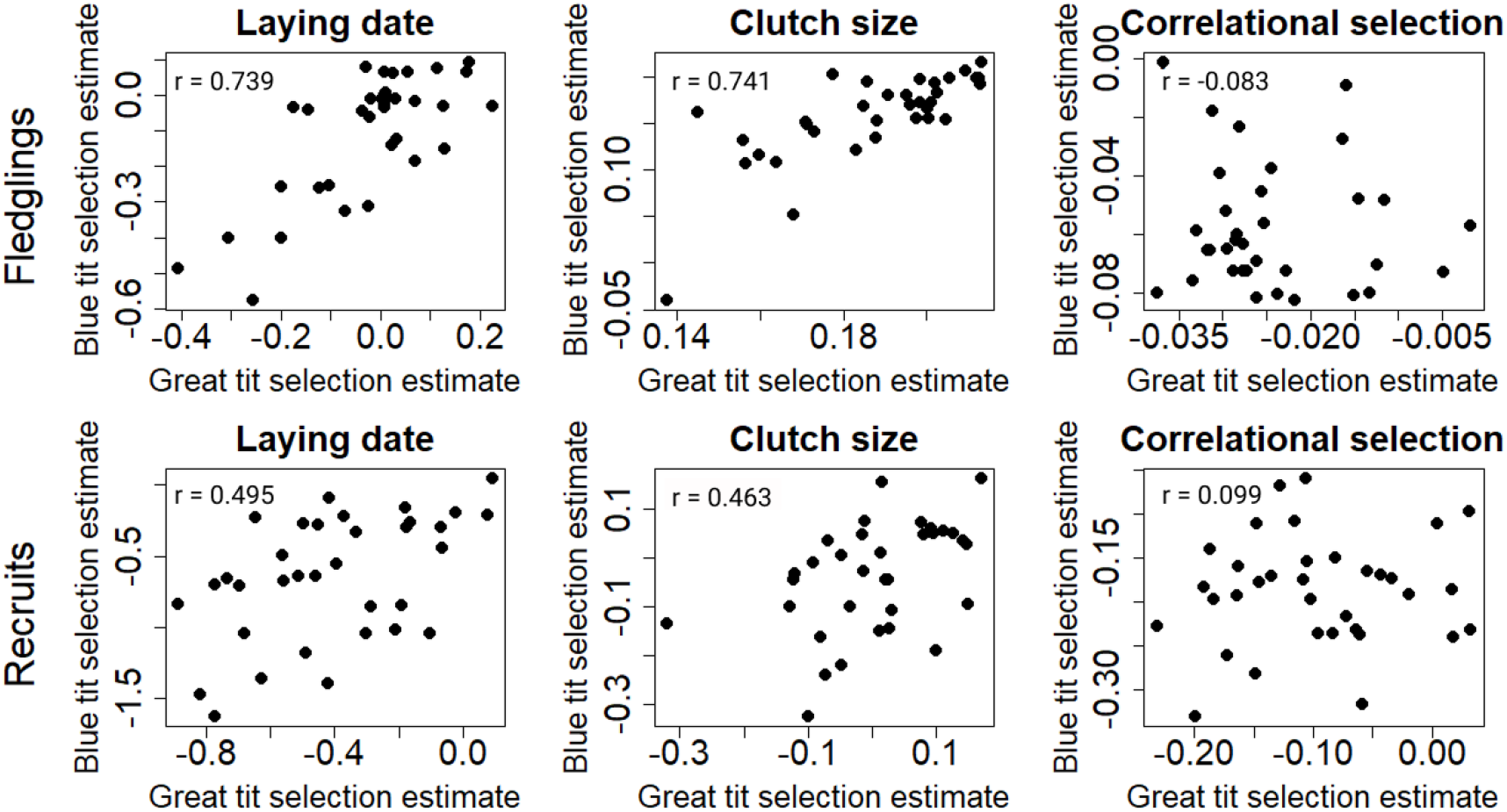
The correlation in selection (with included Pearson correlation value, r) between the great tit and blue tit for the three traits laying date, clutch size and correlational for each year. Data are derived from long-term population study in Wytham Woods, UK, from 1980-2025 (without the years 1991-2000 due to missing fitness data in blue tits. Each point within each plot represents the selection gradient value for both species in a given year for the whole population. In the first row, fledgling numbers are used as fitness measurement while recruit numbers are used at the bottom row.

### Selection regimes in edge-territories

In great tits, we showed that fledgling selection on clutch size tended to be weaker in edge-territories. The same trend was found for recruit selection, although the confidence intervals just overlapped zero. However, this trend was not found for blue tits. We found no evidence for either species to have different selection on laying date between the habitat types. For blue tits, birds breeding in edge-territories produced more dispersers, while no such trend was found for great tits (Table S14). In contrast, great tits with larger clutch sizes produced more recruits that dispersed out of the population (Table S14), which may have led to a small underestimation of selection on clutch size in the recruit model. Otherwise, edge-territories and central territories generally showed similar functions of selection regimes as for the subpopulations (Table 4, S15). Including mother identity in the models provided similar results (Table S15).

**Table 4:** Effect of edge-versus non-edge-territory occupation on selection estimates in great tit and blue tit for clutch size, the quadratic component of clutch size, laying date and the quadratic component of laying date for the models of number of fledglings and number of recruits, respectively. Further is the effect of edge-territories on the fitness variable and the interaction terms with edge-territories and the predictors and their quadratic components. This model is based on a frequentist modelling approach (using the “lme4”-package). The values are the model estimates followed by the 95 % confidence intervals (CI). Further are the random effects including random slope between the random effects and the predictors.

| Species | Great tit |  | Blue tit |  |
| --- | --- | --- | --- | --- |
| Fitness parameter | Fledglings | Recruits | Fledglings | Recruits |
| Fixed effects ($\beta$) | $\beta$ | $\beta$ | $\beta$ | $\beta$ |
| Intercept | 1.81 (1.73, 1.89) | -0.80 (-1.01, -0.62) | 1.88 (1.77, 2.00) | -1.23 (-1.61, -0.82) |
| Clutch size | 0.22 (0.20, 0.23) | 0.10 (0.04, 0.15) | 0.19 (0.16, 0.21) | 0.05 (-0.03, 0.13) |
| Quadratic clutch size | -0.05 (-0.06, -0.04) | -0.08 (-0.11, -0.05) | -0.04 (-0.06, -0.03) | -0.07 (-0.11, -0.03) |
| Laying date | -0.02 (-0.07, 0.03) | -0.43 (-0.54, -0.31) | -0.11 (-0.19, -0.03) | -0.68 (-0.91, -0.44) |
| Quadratic laying date | -0.06 (-0.07, -0.05) | -0.12 (-0.17, -0.06) | -0.05 (-0.07, -0.03) | -0.17 (-0.26, -0.08) |
| Edge-territories | -0.03 (-0.05, 0.00) | -0.15 (-0.23, -0.07) | -0.04 (-0.08, -0.00) | -0.15 (-0.25, -0.05) |
| Altitude (m) | -0.00 (-0.00, -0.00) | 0.00 (0.00, 0.00) | 0.00 (-0.00, 0.00) | -0.00 (-0.00, 0.00) |
| Territory size (ha) | 0.04 (0.03, 0.05) | 0.10 (0.08, 0.12) | 0.01 (-0.00, 0.02) | 0.03 (-0.00, 0.06) |
| Clutch size: Edge-territories | -0.03 (-0.05, -0.01) | -0.06 (-0.13, 0.02) | -0.02 (-0.05, 0.02) | -0.05 (-0.15, 0.05) |
| Quadratic clutch size: Edge-territories | 0.00 (-0.01, 0.02) | 0.00 (-0.04, 0.05) | 0.01 (-0.01, 0.03) | 0.03 (-0.03, 0.10) |
| Laying date: Edge-territories | -0.03 (-0.06, 0.01) | -0.01 (-0.13, 0.09) | -0.05 (-0.10, 0.01) | -0.01 (-0.21, 0.19) |
| Quadratic laying date: Edge-territories | 0.01 (-0.01, 0.02) | 0.02 (-0.06, 0.10) | -0.01 (-0.04, 0.03) | -0.04 (-0.18, 0.10) |
| Random effects ($\sigma$) | $\sigma$ | $\sigma$ | $\sigma$ | $\sigma$ |
| Edge-statusyear: Intercept | 0.046 | 0.098 | 0.268 | 0.000 |
| Edge-statusyear: clutch size | 0.022 | 0.101 | 0.046 | 0.041 |
| Edge-statusyear: laying date | 0.047 | 0.145 | 0.195 | 0.253 |
| Year: Intercept | 0.234 | 0.522 | 0.000 | 1.017 |
| Year: clutch size | 0.021 | 0.057 | 0.000 | 0.113 |
| Year: laying date | 0.144 | 0.298 | 0.004 | 0.509 |
| Sample size (n) | n | n | n | n |
| Number of years | 43 | 43 | 34 | 34 |
| Number of edge-status-years | 86 | 86 | 68 | 68 |
| Number of observations | 10199 | 10105 | 7743 | 7743 |

In general, we found that across fitness traits and species, birds in edge-territories have lower fitness, the exception for fledgling production for great tits (Table 4). In blue tits, this could partly be due to a higher number of dispersers produced in edge-territories (Table S14), while no such effect was found in great tits. While the optimum laying date and clutch size was similar in general over the study period for blue tits (Figure S19 & S20), the optimal clutch size tended to be higher in central territories compared to edge-territories for great tit (Figure S21 & S22).

## Discussion

Here we use a long-term study system to estimate variation in selection across subpopulations and across biologically relevant contrasting habitat types in populations of two ecologically similar species. Selection regimes were similar across contrasting habitats. Most variation in selection among subpopulations was explained by temporal and spatio-temporal sources, depending on the trait examined, while we found little evidence for consistent spatial variation in selection. Further, we found good agreement in selection estimates for the same traits between the two species when comparing spatio-temporal replicates.

### General selection

We show that both species are generally under non-linear selection for larger clutch size and earlier laying date via recruitment and that these traits are under negative correlational selection, when estimated across the entire sample. Similar selection regimes have previously been documented, both in this (only in great tits; (Garant et al. 2007)) and other populations (Porlier et al. 2012; Radchuk et al. 2019; Branston et al. 2021; Jantzen and Visser 2023). Notably, optimum laying dates are closer to the mean for fledgling selection, but significantly earlier than the mean for recruit selection. This is consistent with the idea of scale-dependency, where stabilizing selection is favouring laying dates close to the mean for fledgling success (e.g. representing the scale of the territories the birds experience), while after fledging, the birds experience population-wide processes where hatching early is beneficial (e.g. through increased time for fat storage, density-dependent (Both et al. 1999) and frequency-dependent benefits (Svensson and Connallon 2019)).

### Variation in selection within and between habitats

Our analysis shows clear differences in how selection varied across contexts. The absolute variation in selection was stronger for laying date compared to clutch size and correlational selection, which showed comparable levels of fluctuation. Much of the variation was explained by temporal and spatio-temporal sources. This was largely consistent across traits, fitness parameters and species, although selection on laying date typically had more variation explained by the temporal source than the other traits. In contrast, generally little variation was explained by consistent spatial differences among the subpopulations. The exception was for clutch size for great tits, suggesting that selection for clutch size might have a stronger spatial basis compared to selection for laying date. Notably, this effect was not found for blue tits, which warrants further investigation.

Although the uncertainty intervals of the selection variance components are wide, our results were consistent across traits and species. The results are also supported by frequentist modelling approaches and accounted for sampling error (Morrissey and Hadfield 2012). Temporal variation in selection can also be caused by variance in the mean phenotype among subpopulations (e.g. by low sample size). However, our results are unlikely to be driven by such differences, as standardizing phenotypic traits to the population mean (which reduces the impact of variance in mean phenotypes among the subpopulations) yielded similar results. This is supported by our comparison of variance in the mean phenotypes in general being smaller than the variances in the optimums (Table S13). This is also expected from a theoretical perspective, as environmentally driven changes in the optimum results in a lag between the optimum phenotype and the mean phenotype, creating less variation in the latter, assuming a limited effect of genetic drift, slow evolutionary response and limited adaptive plasticity (Lande and Shannon 1996).

Few studies have decomposed the sources of selection variation within populations (Mouchet et al. 2021; Araya-Ajoy et al. 2025). In our study, the spatio-temporal interaction explained the largest portion of the variance in selection across traits, fitness measure and species. Such spatio-temporal interaction would cover how small-scale spatial variation (e.g. environmental differences among the subpopulations) interact with larger scale temporal variation (e.g. temperature and pressure; Post et al. 1999). This suggests that, when temporal and spatial patterns interact, selection can be highly heterogeneous. While such interactions can maintain genetic diversity, it also has the potential to restrict local adaptation (Nevo 1978), depending on the dispersal capacity of the species. Notably, estimates of direct selection on the same trait in the two species were moderately to strongly correlated at the level of spatio-temporal replicates. This implies that the drivers of selection at this spatio-temporal scale affect each species similarly and hence are unlikely to result from intraspecific mechanisms.

Among populations, previous work has suggested more variation in selection is explained by spatial than temporal sources (Siepielski et al. 2009, 2013, 2017; Morrissey and Hadfield 2012). This is supported by investigations of nearby populations showing that they differ in selection regimes for laying date (Porlier et al. 2012; Jantzen and Visser 2023). However, within a meta-population the spatial component might be expected to be lower. Our finding of limited spatial variation in selection is consistent with studies showing low within-versus among-population variation in selection for behavioural traits (Mouchet et al. 2021) and reproductive traits (Svensson and Sinervo 2004), but differs from previous work in this population highlighting different selection regimes among two subpopulations (Garant et al. 2007). While the spatial effect was lower than expected, it is not surprising because our study system has highly heterogeneous environments (Cole and Sheldon 2017). Tree phenology, an important determinant for laying date (Hinks et al. 2015), is highly variable even at a small scale (Cole and Sheldon 2017). Hence, despite some overall spatial differences (see methods), the subpopulations likely have similar environments in terms of plant phenology. Although the subpopulations varied in density, it can also be possible that there is a consistent scaling of density-dependent selection, although that was not studied here.

The low spatial component is supported by our findings of comparing selection in edge-territories versus central territories, a previously demonstrated important habitat difference (Wilkin et al. 2007a). Edge-territories and central territories were mostly similar selection regimes across species. We found hints of weaker selection for larger clutch size in edge-territories, a trend that was statistically significant for fledgling selection in great tits. In our population, birds breeding in the edges have previously been shown to produce smaller clutches with larger eggs (Wilkin et al. 2007a). The previous study suggested that edge-territories are of poorer quality, favouring larger eggs with higher expected recruitment probability. Our results are partly in line with these findings, as we see signs of weaker selection for clutch size in great tits, although this was not a general result. Indeed, the results mainly imply that the driving force of selection extends equally from central-territories to the edges. Again, this could be driven by our population being highly heterogenous in environmental factors affecting reproductive traits (Hinks et al. 2015; Cole and Sheldon 2017), which could make edge-territories and central territories similar in terms of variation. Interestingly, both species showed reduced fitness in edge-territories, in line with previous studies showing reduced fitness at edges for great tits (Huhta et al. 2004; Wilkin et al. 2007b), although blue tits were not included. Our results on dispersal patterns suggest that lowered fitness in edges is likely due to lower survival, as we only find weak evidence for birds fledged in the edges to have higher dispersal propensity.

More effort has been made to study the temporal variation in selection for these traits (Cresswell and Mccleery 2003; Cao et al. 2019). There are several potential environmental drivers of temporal variation in selection, including temperature (Van Noordwijk et al. 1995), which often affects selection on laying date, potentially via trophic mismatch (Van Noordwijk et al. 1995; Jantzen and Visser 2023). The strong temporal component of selection variation in our population, particularly for laying date, supports the idea that selection fluctuates with large-scale environmental patterns (by changing the optimum phenotype, Visser and Holleman 2001; de Villemereuil et al. 2020) or stochastic demographic effects (by changing population size, density-dependence or mean phenotypes; (Post et al. 1999; Engen and Sæther 2014) that could change among years. Such temporal variation in selection can cause temporal synchrony in selection, where selection across subpopulations co-vary over years, with potential genetic consequences for the population. While fluctuating selection can slow down evolutionary change (Bell 2008; Bonnet and Postma 2018) it can potentially maintain genetic diversity (Nevo 1978; Johnson et al. 2023). However, theoretical models suggest that temporal fluctuating selection is not as efficient as spatially fluctuating selection, which is nearly non-existent here, at maintaining genetic diversity (Nevo 1978). Further, we stress that identifying sources of variation is not equivalent to identifying drivers of selection, calling for more mechanistic studies exploring the causes of selection.

### The role of dispersal

Dispersal is a central concept in studies of varying selection regimes for two reasons. First, dispersal among the subpopulation could affect the phenotypic distribution, and therefore also selection regimes (de Villemereuil et al. 2020). However, the subpopulations showed similar phenotypic distributions. The spatial component for variation in selection is low, reducing the potential impact of dispersal on selection. Further, most individuals in this population breed relatively close to where they were hatched (Greenwood et al. 1979), and breeding dispersal is restricted (Harvey et al. 1979). Hence, we excluded dispersal from our models. Second, dispersal out of the population could affect selection estimation if linked to the studied traits. For great tits, selection estimates could potentially be overestimated (clutch size) and underestimated (quadratic clutch size and correlational selection) as these were associated with dispersal out of Wytham (Table S7). However, no such relationships were found for blue tits. It should be noted, however, that the dispersal dataset was restricted (details in supplementary materials: Dispesal), and these results should be treated with caution.

### Species comparison

Correlation in selection on laying date and clutch size across the two species was high for both fitness traits, but considerably lower when modelling recruit numbers. Although that the two species have shown to be affected by similar environmental drivers in other populations (Van Noordwijk et al. 1995; Porlier et al. 2012), the observed correlation in selection estimates was often surprisingly high, given that at the finest scale studied here, sample sizes were sometimes relatively small, and fitness is often influenced by stochastic variation. In contrast, agreement in correlational selection estimates was low and overlapping zero for both fitness components. Few other studies have explicitly computed the correlation in selection across species, but the correlation would likely vary among traits and the ecological similarity of the species. Such high correlations in selection suggest that the two species have the same, or highly correlated drivers of selection. However, this cannot be concluded in absence of experimental studies. Yet, it is another way to study temporal co-variation in selection. More studies are needed to assess the generality of this finding, both from the same and other species-traits combinations.

## Conclusion

In conclusion, decomposing the variation in selection showed that spatio-temporal interactions govern core drivers of selection. Spatial variation in selection was weak compared to the temporal variation, but the relationship between these sources varied among traits. Such temporal influence on selection could synchronize selection within the population, with potential consequences for genetic diversity and local adaptation. However, spatio-temporal interaction explained considerable variation in selection, suggesting potentially complex genetic and evolutionary impacts. The two species appear to be under similar selection regimes, and differences in central-versus edge-territories were subtle. More studies, particularly from populations with less environmental heterogeneity would be helpful to assess the generality of our findings. Moving forward, identifying the underpinning environmental factors of the spatial, temporal and spatio-temporal sources will be key to understanding how fluctuating selection can affect evolution in the wild.

## Acknowledgement

We would like to thank all the people who have helped to collect data as part of this long-term study over so many years. We would also like to thank Bernt-Erik Sæther and members of the Sheldon-group for feedback on this project.

## Author contributions

JSS, YAA, EFC and BCS conceived the study. JSS, EFC and BCS collected data and JSS analysed the data with input from all authors. JSS wrote the manuscript with input from all authors.

