## Supplementaryt materials for "Fine-scale spatio-temporal variation in natural selection on reproductive traits across two wild bird species"

**Supplementary materials**

***Modelling of variation in selection in subpopulations in Wytham***

We modelled the fitness (either number of fledglings or number of recruits produced) of individual m in subpopulation i in year j and subpopulation-year k as the following;

**Equation 1:**

V_ijkm_ = ß_0_ + ß_a_ + ß_b_ + ß_c_ **+** ß_d_+ ß_e_+ µ_0i_ + µ_1i_ + µ_2i_ + µ_3i_ + µ_4i_ + µ_5i_ + Ʊ_0k_ + Ʊ_1k_ + Ʊ_2k_ + Ʊ_3k_ + Ʊ_4k_ + Ʊ_5k_ + ν_0j_ + ν_1j_ + ν_2j_ + ν_3j_ + ν_4j_ + ν_5j_

where V is an individual’s expected log fitness *ß_0_* is the intercept of the model, along with the following fixed effects; *ßa* is the effect of clutch size, *ß_b_* is the effect of quadratic clutch size, *ß_c_* is the effect of laying date , *ß_d_* is the effect of quadratic laying date and *ß_e_* represent the interaction between laying date and clutch size. All the levels of random effects, that is, subpopulations µ_i_, years ν_j_ and subpopulation-year Ʊ_k_ allowed for random intercept (0), random slope for clutch size (1), random slope for quadratic clutch size (2), random slope for laying date (3), random slope for quadratic laying date (4) and random slope for the interaction between laying date and clutch size (3).

***Modelling of selection in edge-territories versus central territories***

We modelled the selection regimes in edge-territories versus central territories by modelling fitness (either number of fledglings or number of recruits produced) of individual m in edge-status-year i and year j as the following;

**Equation 2:**

V_ijkm_ = ß_0_ + ß_a_ a + ß_b_ b + ß_c_ c **+** ß_d_ d+ ß_e_ e+ ß_f_ f+ ß_g_ g+ ß_h_ h + ß_i_ i + ß_j_ j + ß_k_ k + µ_0i_ + µ_1i_ + µ_2i_ + ν_0j_ + ν_1j_ + ν_2j_

where v is an individual’s expected log fitness *ß_0_* is the intercept of the model, along with the following fixed effects; *ß_a_* is the effect of clutch size (a), *ß_b_* is the effect of quadratic clutch size (b), *ß_c_* is the effect of laying date (c) and *ß_d_* is the effect of quadratic laying date (d). To investigate if these patterns differed between edges and non-edges, we also included *ß_e_*, which is the interaction between edge-status and clutch size (e), *ß_f_* representing the interaction between edge-status and quadratic clutch size (f), *ß_g_* the interaction between edge-status and laying date (g) and *ß_h_,* representing the interaction between edge-status and quadratic laying date (h). Further, ß_i_ represents the effect of altitude (i), ß_j_ represents the effect of territory size (j) and ß_k_ k represents the effect of being in an edge-territory (binary variable (1/0)). All the levels of random effects, that is, edge-status-year *µ_i_* and years *ν_j_*, allowed for random intercept (0), random slope for clutch size (1) and random slope for laying date (2).

***Testing for biases in selection***

To check if the proportion of variation in selection was driven by differences in mean phenotypes among the subpopulations rather than changes in the optimum, we also performed the same model as above, but standardizing each trait to the population mean rather than the subpopulation mean. This reduces the variation in mean phenotype both spatially and temporally. We also checked if female identity could affect our results by including this as a random effect in an additional model otherwise structured as the models in the main text. Initially, all models were run without female identity, as females with earlier laying dates were less likely to be detected. Therefore, including only identified females would bias our estimates of laying date. Further, we also tested how fledgling numbers affected the number of recruits produced by using the same model structure as described above but also including fledgling numbers as fixed effect when modelling recruit numbers. To investigate how laying date affected clutch size, we used a linear mixed model with standardized clutch size as a response variable and standardized laying date and its quadratic component as fixed effects. Subpopulation, year and subpopulation-year were used as random effects. When using the number of recruits as fitness measurement, selection estimates could be biased if the trait of interest co-varies with the number of birds that disperse out of the population. To investigate if such dispersal could affect our selection estimate, we used a small subset of known recoveries outside of our population and modelled the number of dispersers by the same fixed effects as above. In addition, we included the same random effects, but did not allow for random slopes due to low sample sizes.

**Supplementary figures**

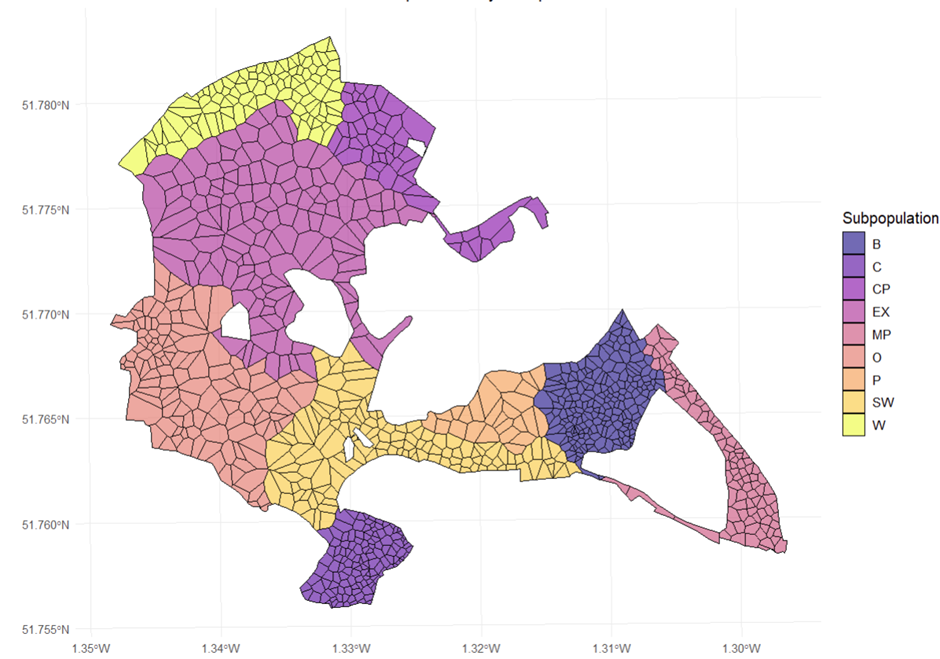

**Figure S1:** The different subpopulations of Wytham marked in different colours. The lines included in the spatial plot represent the territory boundaries if all nest-boxes are occupied. Subpopulation P was excluded from the analysis due to consistent low sample size.

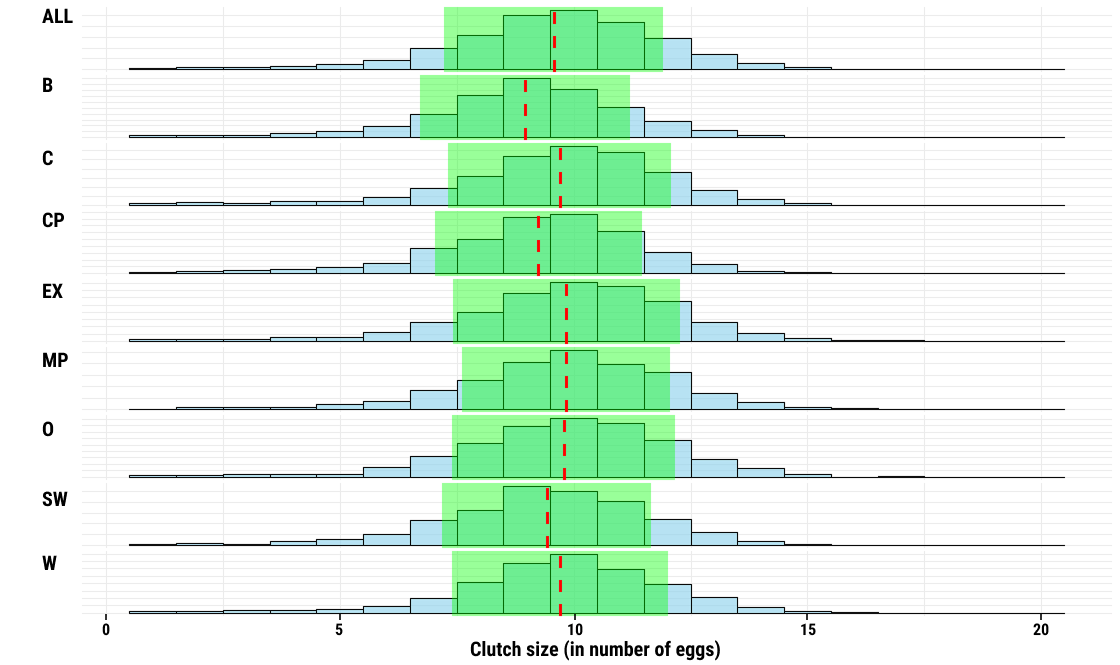

**Figure S2:** Distribution of clutch size for the entire population (top) and all the subpopulations for blue tits. The red lines represent the (sub) population mean and the green encompass its standard deviation.

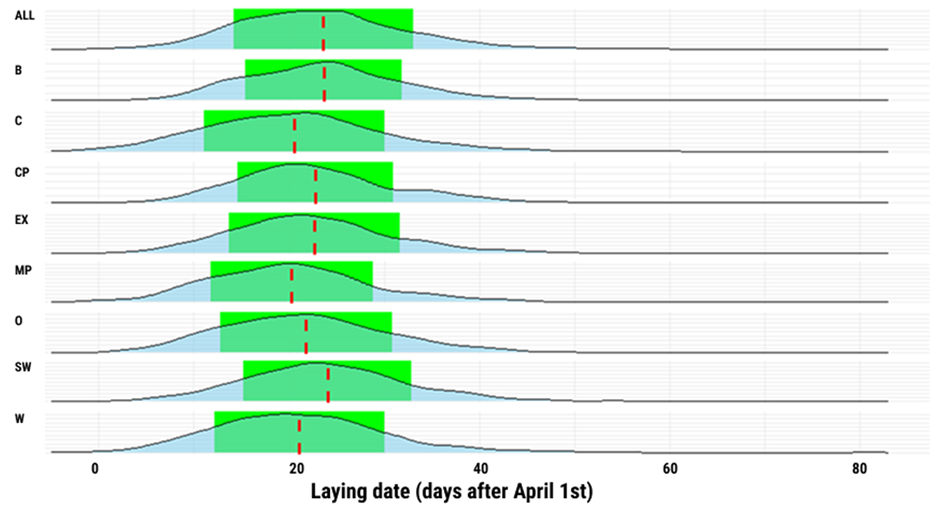

**Figure S3:** Distribution of laying date (in days after April 1st) for the entire population (top) and all the subpopulations for blue tits. The red lines represent the (sub)population mean and the green encompass its standard deviation.

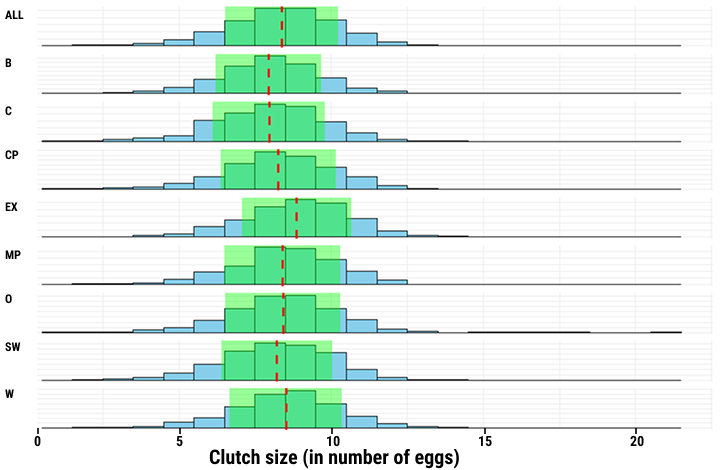
**Figure S4**: Overview over clutch size for the entire population (top) and all the subpopulations for great tits. The red lines represent the (sub)population’s mean and the green encompass its standard deviation.

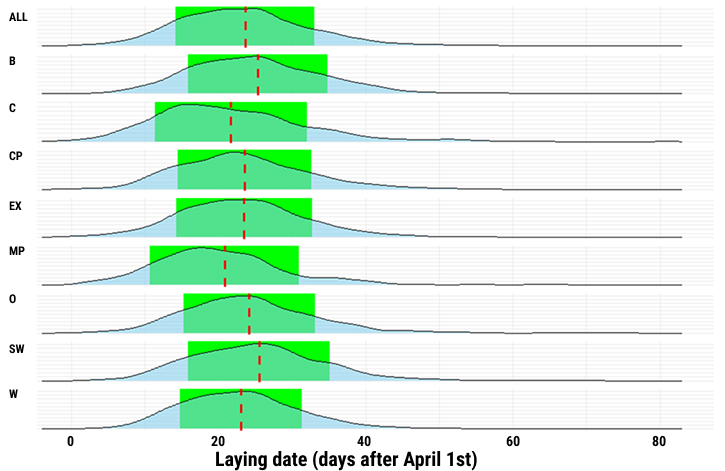

**Figure S5**: Overview over laying date (in days after April 1st) for the entire population (top) and all the subpopulations for great tits. The red lines represent the (sub)population’s mean and the green encompass its standard deviation.

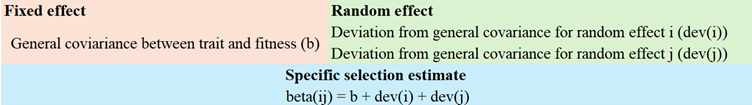

**Figure S6:** Visual representation of how selection estimates for a particular unit (beta(ij)) can be extracted using the method described in the methods-section. The general selection estimate can be combined by the deviation from the general covariance caused by each different random effect. If more than one random effect is relevant for the specific unit, these are added as well.

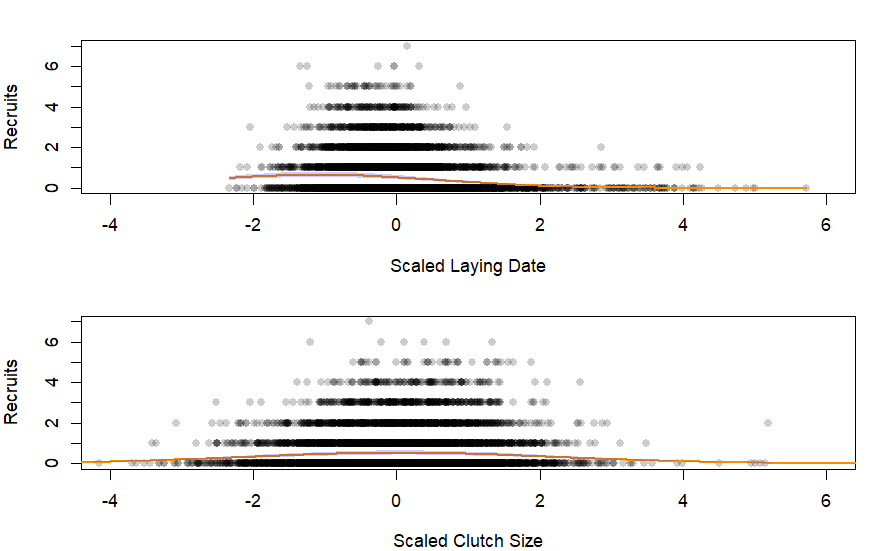

**Figure S7**: The relationship between scaled laying date and the number of recruits produced in the top figure and between scaled clutch size and the number of recruits produced for great tit. The shaded area represents the CI of the estimated lines. Each circle represents a breeding attempt, and these circles are transparent to highlight overlapping data points.

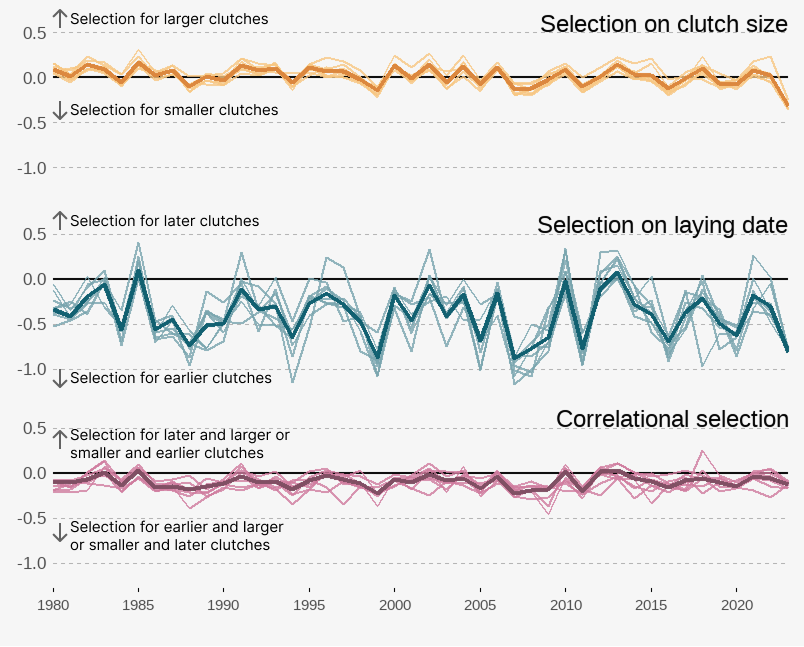

**Figure S8:** Yearly overview of the selection estimates for clutch size, laying date and the correlational selection coefficients between the two for great tit. Each line with the weaker color represents a subpopulation while the line with stronger color represents the population-wide selection estimate. The selection estimates are extracted from a frequentist modelling approach (using the “lme4”-package). To extract the selection estimate for a given subpopulation for a given year, we combined the model estimate for the trait of interest with the estimate for that trait's random slope of year, subpopulation and subpopulation-year. By adding these together, one obtains the overall selection estimate for each subpopulation for each year. The population-wide estimate was extracted by combining the model estimate for the trait of interest with the estimate for that trait's random slope of year. The limit of the y-axis is the same for all three panels to facilitate comparison of magnitude of variation in selection among the traits.

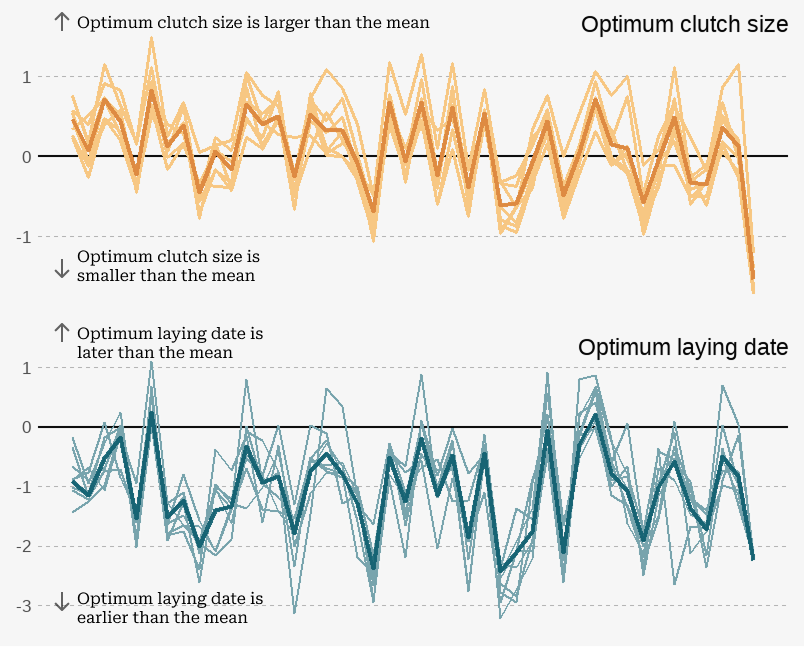

**Figure S9:** Yearly overview over the estimated optimal phenotype for recruit production in each of the different subpopulations (weakly colored lines) and for the entire population (strongly colored line) for great tit. The optimum is calculated as the linear selection estimate over two times the negative estimate of the quadratic (non-linear selection) component. The selection estimates are extracted from a frequentist modelling approach (using the “lme4”-package). To extract the selection estimate for a given subpopulation for a given year, we combined the model estimate for the trait of interest with the estimate for that trait's random slope of year, subpopulation and subpopulation-year. By adding these together, one obtains the overall selection estimate for each subpopulation for each year. The population-wide estimate was extracted by combining the model estimate for the trait of interest with the estimate for that trait's random slope of year.

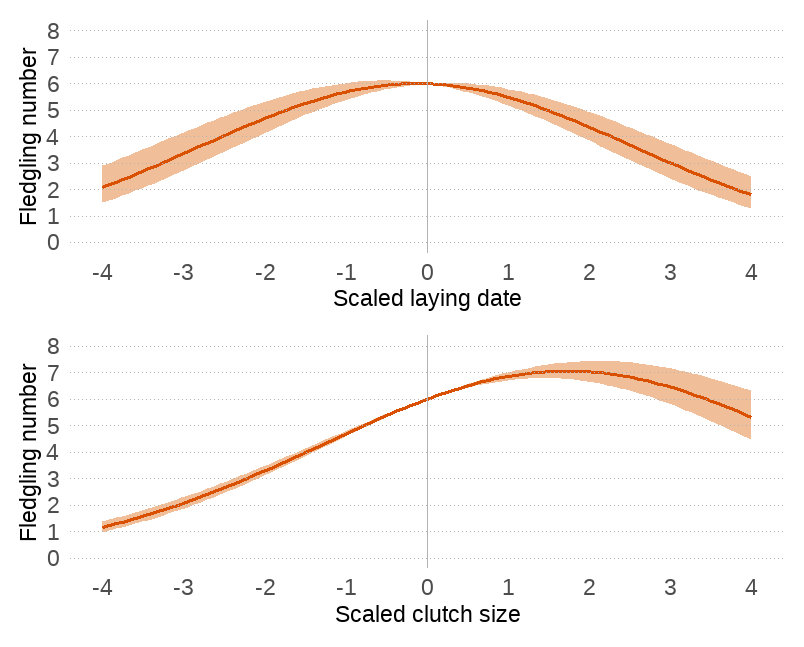

**Figure S10:** The relationship between a) scaled laying date and the number of fledglings produced and b) between scaled clutch size and the number of fledglings produced for great tit. Each trait is measured in the number of standard deviations from the mean. The shaded area represents the CI of the estimated lines. Data points are excluded from this figure to facilitate interpretation (see Figure S7 for data points included).

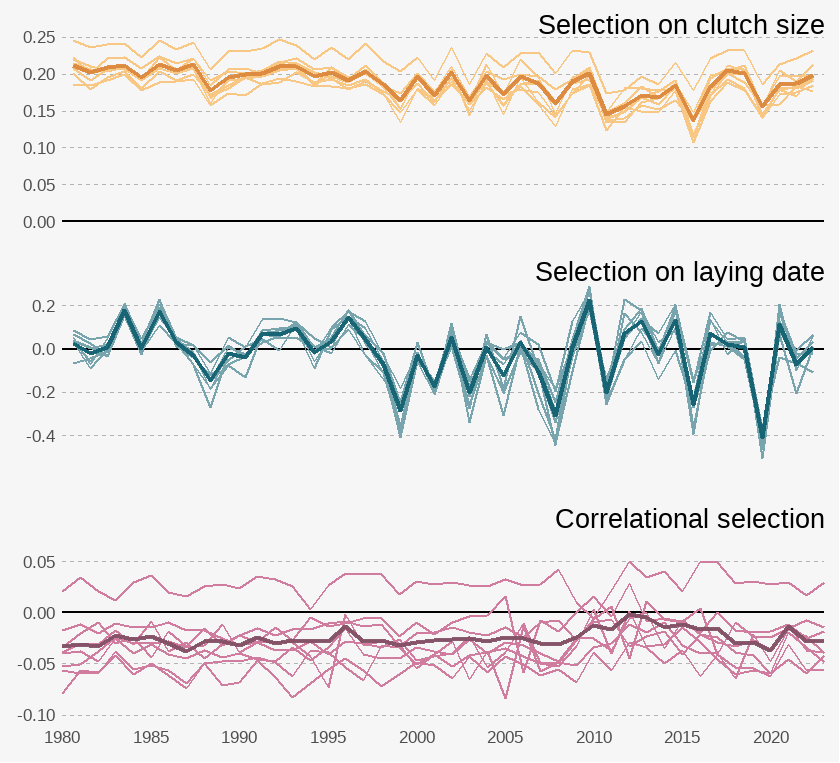

**Figure S11:** Yearly overview of the selection estimates for clutch size, laying date and the correlational selection coefficients between the two for great tit when treating fledgling numbers as fitness. Each line with the weaker color represents a subpopulation while the line with stronger color represents the population-wide selection estimates. The selection estimates are extracted from a frequentist modelling approach (using the “lme4”-package). To extract the selection estimate for a given subpopulation for a given year, we combined the model estimate for the trait of interest with the estimate for that trait's random slope of year, subpopulation and subpopulation-year. By adding these together, one obtains the overall selection estimate for each subpopulation for each year. The population-wide estimate was extracted by combining the model estimate for the trait of interest with the estimate for that trait's random slope of year.

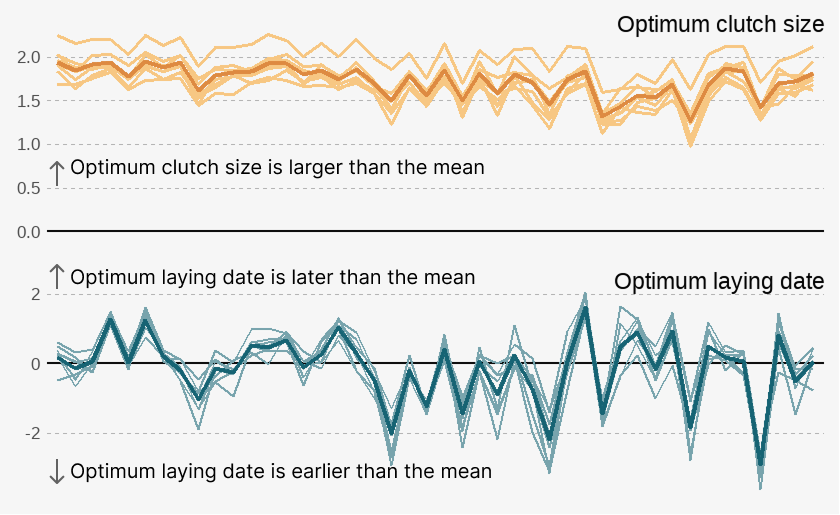

**Figure S12:** Yearly overview of the estimated optimal phenotype for fledgling production in each of the different subpopulations (weakly colored lines) and for the entire population (strongly colored line) for great tit. The optimum is calculated as the linear selection estimate over two times the negative estimate of the quadratic (non-linear selection) component. The selection estimates are extracted from a frequentist modelling approach (using the “lme4”-package). To extract the selection estimate for a given subpopulation for a given year, we combined the model estimate for the trait of interest with the estimate for that trait's random slope of year, subpopulation and subpopulation-year. By adding these together, one obtains the overall selection estimate for each subpopulation for each year. The population-wide estimate was extracted by combining the model estimate for the trait of interest with the estimate for that trait's random slope of year. The limit of the y-axis is the same for all three panels to facilitate comparison of magnitude of variation in selection among the traits.

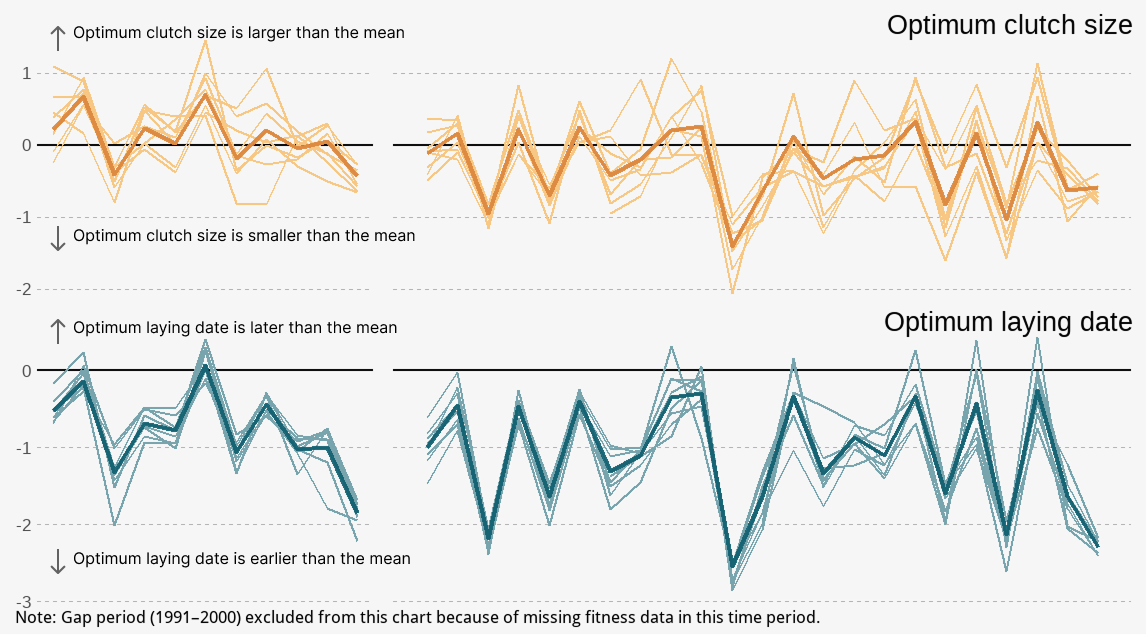

**Figure S13:** Yearly overview over the estimated optimal phenotype for recruit production in each of the different subpopulations (weakly colored lines) and for the entire population (strongly colored line) for blue tit. The optimum is calculated as the linear selection estimate over two times the negative estimate of the quadratic (non-linear selection) component. The selection estimates are extracted from a frequentist modelling approach (using the “lme4”-package). To extract the selection estimate for a given subpopulation for a given year, we combined the model estimate for the trait of interest with the estimate for that trait's random slope of year, subpopulation and subpopulation-year. By adding these together, one obtains the overall selection estimate for each subpopulation for each year. The population-wide estimate was extracted by combining the model estimate for the trait of interest with the estimate for that trait's random slope of year. The limit of the y-axis is the same for all three panels to facilitate comparison of magnitude of variation in selection among the traits.

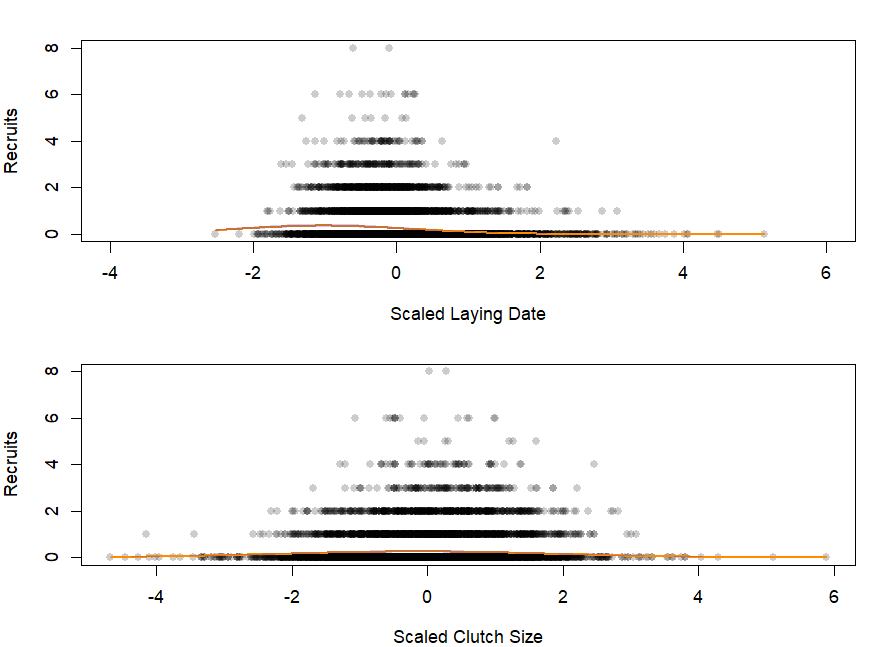

**Figure S14:** The relationship between scaled laying date and the number of recruits produced in the top figure and between scaled clutch size and the number of recruits produced for blue tit The shaded area represents the CI of the estimated lines. Each circle represents a breeding attempt, and these circles are transparent to highlight overlapping data points.

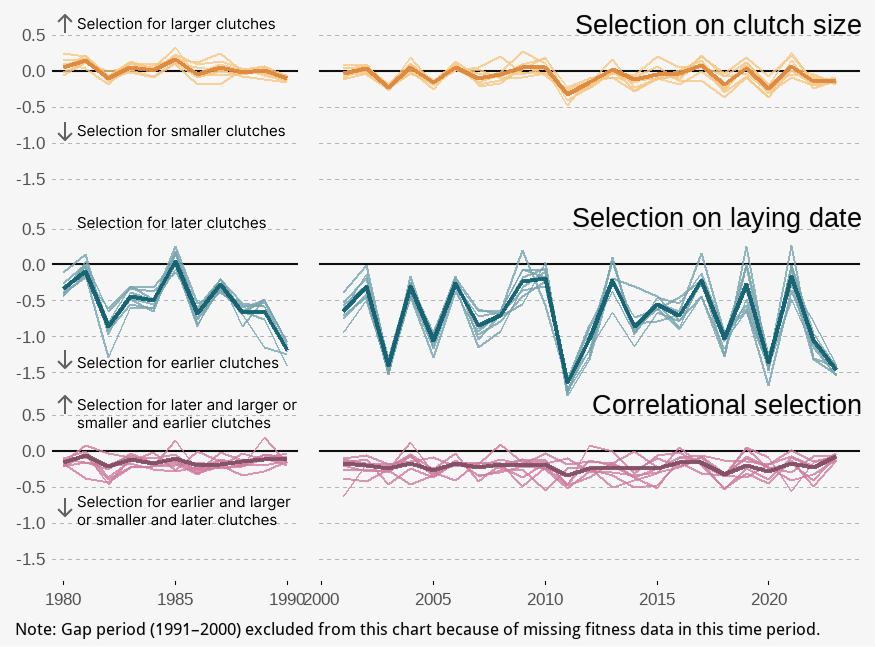

**Figure S15:** Yearly overview of the selection estimates for clutch size, laying date and the correlational selection coefficients between the two for blue tit. Each line with the weaker color represents a subpopulation while the line with stronger color represents the population-wide selection estimates. The selection estimates are extracted from a frequentist modelling approach (using the “lme4”-package). To extract the selection estimate for a given subpopulation for a given year, we combined the model estimate for the trait of interest with the estimate for that trait's random slope of year, subpopulation and subpopulation-year. By adding these together, one obtains the overall selection estimate for each subpopulation for each year. The population-wide estimate was extracted by combining the model estimate for the trait of interest with the estimate for that trait's random slope of year. The limit of the y-axis is the same for all three panels to facilitate comparison of magnitude of variation in selection among the traits.

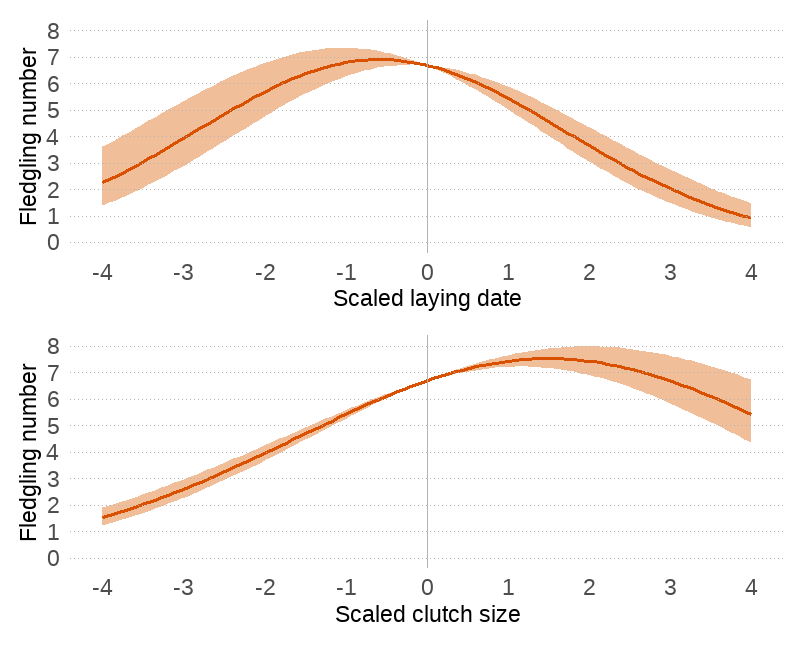

**Figure S16:** The relationship between a) scaled laying date and the number of fledglings produced and b) between scaled clutch size and the number of fledglings produced for blue tit. Each trait is measured in the number of standard deviations from the mean. The shaded area represents the CI of the estimated lines. Data points are excluded from this figure to facilitate interpretation (see Figure S14 for data points included).

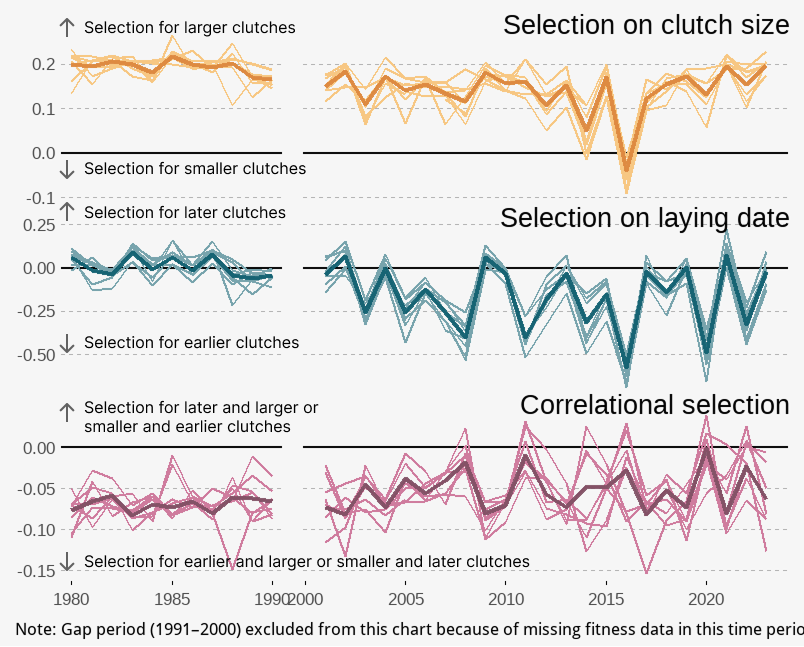

**Figure S17:** Yearly overview of the selection estimates for clutch size, laying date and the correlational selection coefficients between the two for blue tit when treating fledgling numbers as fitness. Each line with the weaker color represents a subpopulation while the line with stronger color represents the population-wide selection estimates. The selection estimates are extracted from a frequentist modelling approach (using the “lme4”-package). To extract the selection estimate for a given subpopulation for a given year, we combined the model estimate for the trait of interest with the estimate for that trait's random slope of year, subpopulation and subpopulation-year. By adding these together, one obtains the overall selection estimate for each subpopulation for each year. The population-wide estimate was extracted by combining the model estimate for the trait of interest with the estimate for that trait's random slope of year.

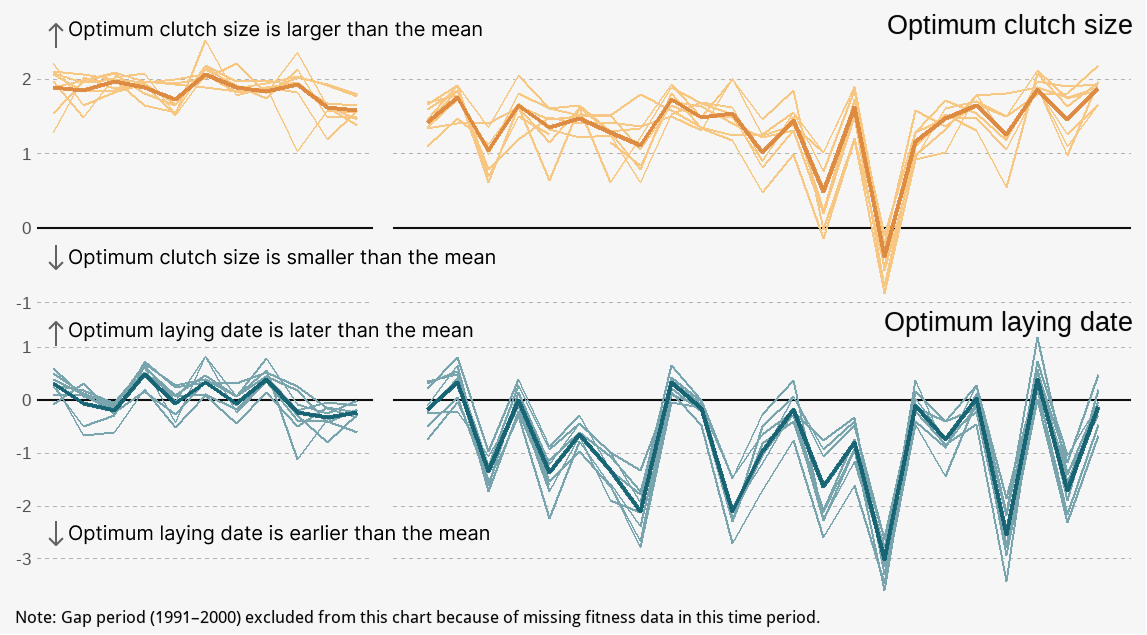

**Figure S18:** Yearly overview over the estimated optimal phenotype for fledgling production in each of the different subpopulations (weakly colored lines) and for the entire population (strongly colored line) for blue tit. The optimum is calculated as the linear selection estimate over two times the negative estimate of the quadratic (non-linear selection) component. The selection estimates are extracted from a frequentist modelling approach (using the “lme4”-package). To extract the selection estimate for a given subpopulation for a given year, we combined the model estimate for the trait of interest with the estimate for that trait's random slope of year, subpopulation and subpopulation-year. By adding these together, one obtains the overall selection estimate for each subpopulation for each year. The population-wide estimate was extracted by combining the model estimate for the trait of interest with the estimate for that trait's random slope of year. The limit of the y-axis is the same for all three panels to facilitate comparison of magnitude of variation in selection among the traits.

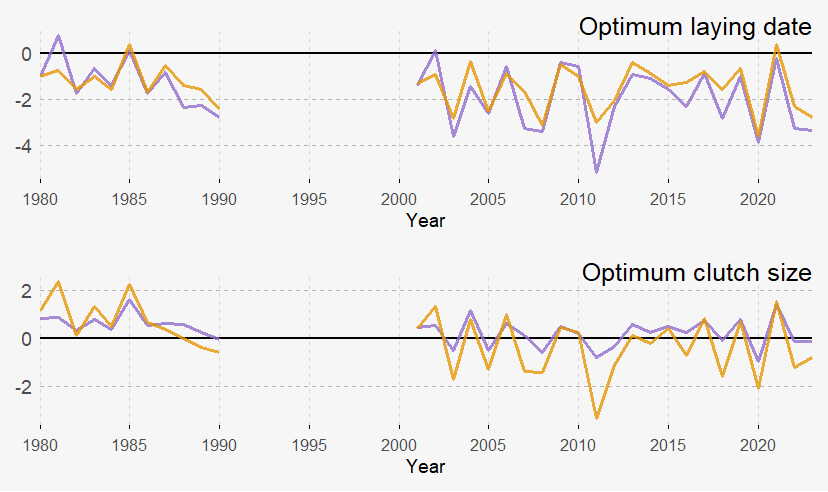

**Figure S19:** The optimal clutch size and laying date for central,- and edge-territories over the study period for blue tit when the number of recruits is considered as fitness. The optimum is calculated as the linear selection estimate over two times the negative estimate of the quadratic (non-linear selection) component. The selection estimates are extracted from a frequentist modelling approach (using the “lme4”-package). To extract the selection estimate for a given edge-status for a given year, we combined the model estimate for the trait of interest with the estimate for that trait's random slope of year and edge-status year. By adding these together, one obtains the overall selection estimate for each edge-status for each year. The years 1991-2000 are excluded due to missing fitness data in these years. Central territories are marked in lilac and edge-territories are marked in orange.

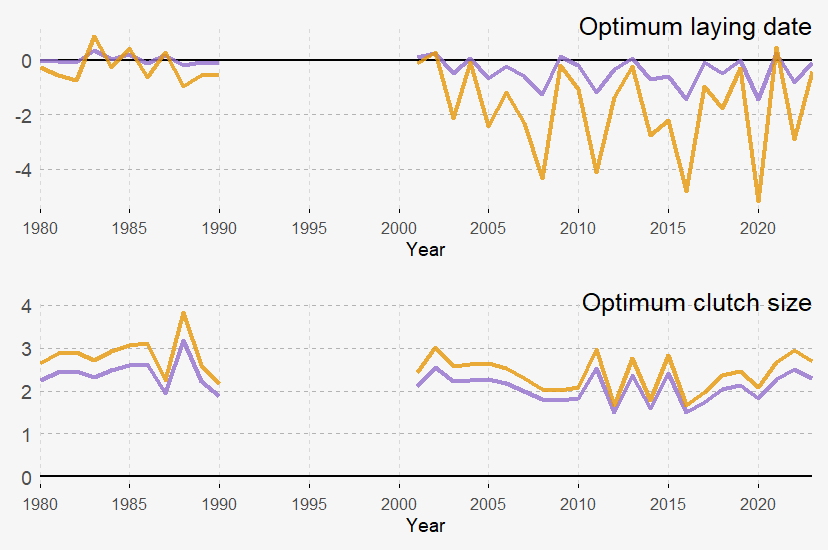

**Figure S20:** The optimal clutch size and laying date for central,- and edge-territories over the study period for blue tit when the number of fledglings is considered as fitness. The optimum is calculated as the linear selection estimate over two times the negative estimate of the quadratic (non-linear selection) component. The selection estimates are extracted from a frequentist modelling approach (using the “lme4”-package). To extract the selection estimate for a given edge-status for a given year, we combined the model estimate for the trait of interest with the estimate for that trait's random slope of year and edge-status year. By adding these together, one obtains the overall selection estimate for each edge-status for each year. The years 1991-2000 are excluded due to missing fitness data in these years. Central territories are marked in lilac and edge-territories are marked in orange

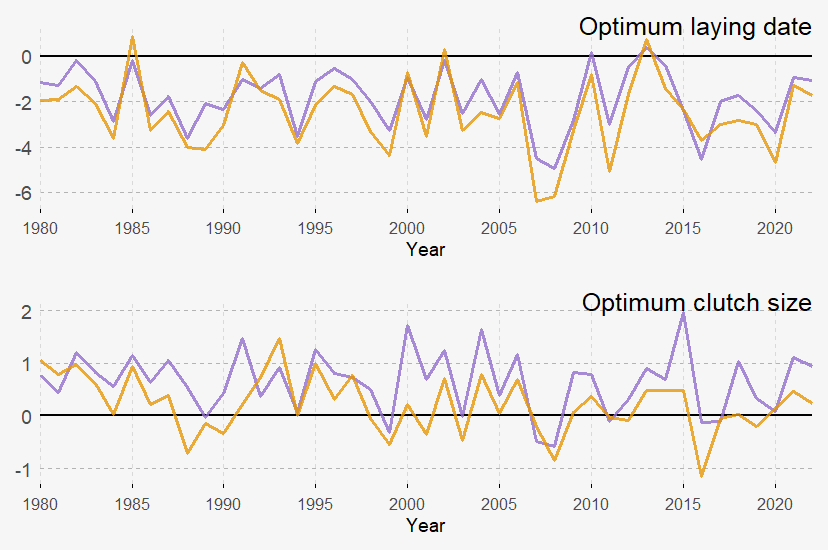

**Figure S21:** The optimal clutch size and laying date for central,- and edge-territories over the study period for great tit when the number of recruits is considered as fitness. The optimum is calculated as the linear selection estimate over two times the negative estimate of the quadratic (non-linear selection) component. The selection estimates are extracted from a frequentist modelling approach (using the “lme4”-package). To extract the selection estimate for a given edge-status for a given year, we combined the model estimate for the trait of interest with the estimate for that trait's random slope of year and edge-status year. By adding these together, one obtains the overall selection estimate for each edge-status for each year. Central territories are marked in lilac and edge-territories are marked in orange.

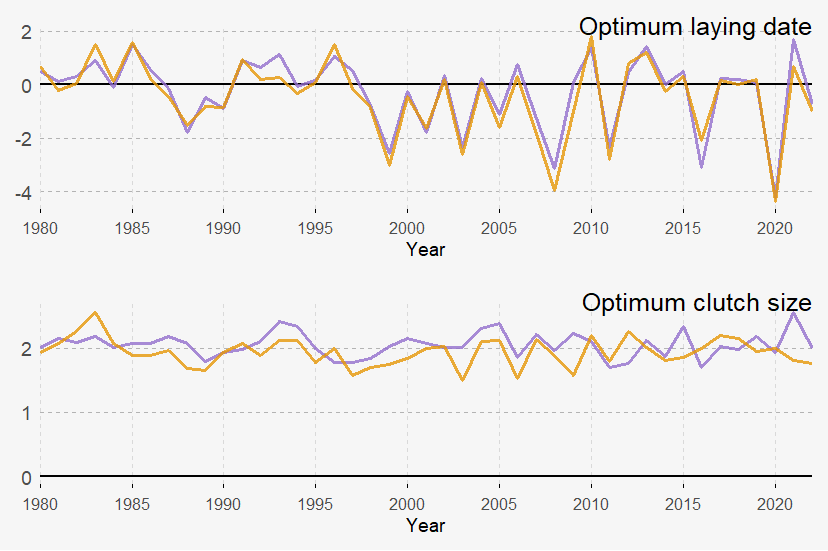

**Figure S22:** The optimal clutch size and laying date for central,- and edge-territories over the study period for great tit when the number of fledglings is considered as fitness. The optimum is calculated as the linear selection estimate over two times the negative estimate of the quadratic (non-linear selection) component. The selection estimates are extracted from a frequentist modelling approach (using the “lme4”-package). To extract the selection estimate for a given edge-status for a given year, we combined the model estimate for the trait of interest with the estimate for that trait's random slope of year and edge-status year. By adding these together, one obtains the overall selection estimate for each edge-status for each year. Central territories are marked in lilac and edge-territories are marked in orange.

**SUPPLEMENTARY TABLES**

**Table S1:** Yearly summary of the number of reproductive events of great tits per subpopulation per year that could be included in this study. Clutches subject to experimental manipulations are excluded from the study and the table, leading to lower sample sizes in certain years and certain subpopulations.

| **Year** | **b** | **c** | **cp** | **ex** | **mp** | **o** | **sw** | **w** |
| --- | --- | --- | --- | --- | --- | --- | --- | --- |
| 1980 | 16 | 17 | 8 | 29 | 12 | 24 | 23 | 14 |
| 1981 | 42 | 32 | 17 | 67 | 26 | 37 | 41 | 32 |
| 1982 | 29 | 21 | 14 | 49 | 6 | 29 | 29 | 20 |
| 1983 | 46 | 28 | 25 | 83 | 6 | 55 | 53 | 36 |
| 1984 | 36 | 14 | 12 | 50 | 3 | 31 | 29 | 37 |
| 1985 | 30 | 10 | 12 | 40 | 10 | 28 | 26 | 24 |
| 1986 | 34 | 21 | 14 | 40 | 14 | 27 | 27 | 24 |
| 1987 | 33 | 19 | 15 | 48 | 26 | 32 | 32 | 29 |
| 1988 | 46 | 23 | 18 | 55 | 31 | 35 | 28 | 35 |
| 1989 | 26 | 21 | 18 | 48 | 25 | 29 | 14 | 32 |
| 1990 | 33 | 24 | 12 | 41 | 3 | 16 | 12 | 27 |
| 1991 | 17 | 18 | 5 | 12 | 10 | 14 | 19 | 13 |
| 1992 | 22 | 16 | 17 | 44 | 5 | 30 | 21 | 34 |
| 1993 | 46 | 25 | 25 | 56 | 36 | 36 | 43 | 43 |
| 1994 | 43 | 26 | 19 | 57 | 28 | 49 | 45 | 48 |
| 1995 | 38 | 25 | 25 | 74 | 29 | 59 | 39 | 38 |
| 1996 | 46 | 31 | 29 | 88 | 35 | 52 | 40 | 54 |
| 1997 | 43 | 29 | 25 | 68 | 34 | 48 | 36 | 40 |
| 1998 | 33 | 25 | 23 | 77 | 37 | 38 | 44 | 48 |
| 1999 | 33 | 24 | 16 | 50 | 37 | 34 | 28 | 34 |
| 2000 | 28 | 38 | 16 | 39 | 33 | 27 | 35 | 33 |
| 2001 | 46 | 36 | 11 | 41 | 16 | 12 | 26 | 25 |
| 2002 | 51 | 44 | 16 | 35 | 15 | 9 | 24 | 25 |
| 2003 | 57 | 39 | 28 | 59 | 50 | 38 | 61 | 43 |
| 2004 | 19 | 33 | 11 | 41 | 20 | 35 | 25 | 36 |
| 2005 | 67 | 56 | 32 | 79 | 44 | 62 | 69 | 52 |
| 2006 | 46 | 37 | 28 | 69 | 46 | 39 | 35 | 56 |
| 2007 | 51 | 41 | 29 | 85 | 51 | 38 | 40 | 73 |
| 2008 | 53 | 29 | 18 | 67 | 46 | 48 | 60 | 53 |
| 2009 | 12 | 36 | 15 | 40 | 14 | 25 | 27 | 28 |
| 2010 | 47 | 35 | 24 | 65 | 39 | 44 | 47 | 56 |
| 2011 | 26 | 22 | 14 | 26 | 22 | 27 | 27 | 21 |
| 2012 | 49 | 42 | 24 | 71 | 42 | 50 | 60 | 44 |
| 2013 | 41 | 29 | 16 | 41 | 31 | 31 | 38 | 31 |
| 2014 | 22 | 17 | 9 | 26 | 23 | 14 | 31 | 12 |
| 2015 | 47 | 29 | 21 | 57 | 41 | 41 | 56 | 40 |
| 2016 | 30 | 16 | 18 | 60 | 39 | 34 | 42 | 37 |
| 2017 | 39 | 19 | 19 | 62 | 23 | 45 | 50 | 39 |
| 2018 | 26 | 22 | 22 | 57 | 23 | 48 | 41 | 33 |
| 2019 | 44 | 32 | 22 | 73 | 38 | 54 | 58 | 54 |
| 2020 | 35 | 25 | 16 | 36 | 33 | 34 | 37 | 36 |
| 2021 | 32 | 25 | 19 | 38 | 32 | 34 | 39 | 39 |
| 2022 | 32 | 30 | 18 | 41 | 23 | 34 | 36 | 33 |
| 2023 | 29 | 20 | 17 | 51 | 29 | 33 | 40 | 34 |

**Table S2:** Yearly summary of the number of reproductive events of blue tits per subpopulation per year that could be included in this study. Clutches subject to experimental manipulations are excluded from the study and the table, leading to lower sample sizes in certain years and certain subpopulations.

| **year** | **b** | **c** | **cp** | **ex** | **mp** | **o** | **sw** | **w** |
| --- | --- | --- | --- | --- | --- | --- | --- | --- |
| 1980 | 43 | 27 | 8 | 32 | 36 | 45 | 45 | 27 |
| 1981 | 61 | 47 | 17 | 30 | 44 | 49 | 45 | 53 |
| 1982 | 60 | 33 | 19 | 34 | 5 | 41 | 30 | 43 |
| 1983 | 52 | 40 | 11 | 25 | 8 | 31 | 43 | 43 |
| 1984 | 47 | 44 | 13 | 32 | 2 | 43 | 38 | 33 |
| 1985 | 50 | 50 | 20 | 54 | 22 | 46 | 39 | 58 |
| 1986 | 39 | 45 | 15 | 29 | 22 | 31 | 16 | 43 |
| 1987 | 44 | 43 | 12 | 24 | 23 | 9 | 12 | 16 |
| 1988 | 61 | 44 | 8 | 22 | 36 | 23 | 29 | 53 |
| 1989 | 56 | 45 | 9 | 25 | 35 | 54 | 34 | 12 |
| 1990 | 70 | 57 | 11 | 65 | 29 | 59 | 47 | 44 |
| 2001 | 16 | 19 | 4 | 17 | 13 | 20 | 13 | 14 |
| 2002 | 14 | 7 | 6 | 15 | 6 | 17 | 14 | 13 |
| 2003 | 13 | 21 | 11 | 20 | 12 | 26 | 10 | 9 |
| 2004 | 49 | 27 | 29 | 20 | 27 | 40 | 25 | 25 |
| 2005 | 43 | 22 | 24 | 30 | 26 | 47 | 27 | 22 |
| 2006 | 44 | 4 | 4 | 8 | 28 | 32 | 34 | 2 |
| 2007 | 56 | 20 | 18 | 22 | 24 | 51 | 55 | 6 |
| 2008 | 54 | 40 | 24 | 16 | 29 | 5 | 42 | 11 |
| 2009 | 50 | 35 | 17 | 38 | 37 | 27 | 32 | 17 |
| 2010 | 74 | 50 | 49 | 74 | 33 | 79 | 59 | 44 |
| 2011 | 91 | 62 | 25 | 72 | 48 | 88 | 73 | 48 |
| 2012 | 65 | 44 | 44 | 72 | 44 | 81 | 49 | 58 |
| 2013 | 40 | 21 | 33 | 61 | 32 | 61 | 32 | 44 |
| 2014 | 45 | 42 | 42 | 68 | 34 | 83 | 48 | 52 |
| 2015 | 40 | 28 | 11 | 40 | 22 | 67 | 33 | 43 |
| 2016 | 42 | 42 | 44 | 67 | 21 | 81 | 57 | 49 |
| 2017 | 24 | 31 | 35 | 58 | 29 | 67 | 33 | 51 |
| 2018 | 37 | 41 | 38 | 61 | 30 | 72 | 38 | 54 |
| 2019 | 47 | 50 | 54 | 71 | 36 | 85 | 55 | 53 |
| 2020 | 77 | 65 | 55 | 108 | 52 | 96 | 78 | 74 |
| 2021 | 60 | 64 | 54 | 107 | 46 | 102 | 59 | 65 |
| 2022 | 57 | 52 | 50 | 96 | 25 | 98 | 63 | 68 |
| 2023 | 30 | 46 | 44 | 78 | 30 | 92 | 68 | 63 |

**Table S3:** Distribution of different habitat types among the different subpopulations in Wytham. Included is also information on the mean (in meters) and standard deviation of altitude of the nestboxes in the different subpopulations.

|  | **B** | **C** | **CP** | **EX** | **MP** | **O** | **SW** | **W** |
| --- | --- | --- | --- | --- | --- | --- | --- | --- |
| **Proportion of boxes in habitats** |  |  |  |  |  |  |  |  |
| Twentieth century plantation | 0.02 | 0 | 0 | 0.19 | 0 | 0.57 | 0.31 | 0 |
| Eighteen and nineteen century plantation | 0.04 | 0 | 0.18 | 0.20 | 1 | 0 | 0.55 | 0 |
| Semi-natural woodland | 0.94 | 1 | 0 | 0.41 | 0 | 0.20 | 0.13 | 1 |
| Secondary woodland | 0 | 0 | 0.82 | 0.20 | 0 | 0.23 | 0.006 | 0 |
| **Altitude information** |  |  |  |  |  |  |  |  |
| Mean altitude (m) | 110.8 | 99.8 | 94.1 | 131.5 | 73.2 | 105.1 | 132.7 | 72.3 |
| Standard deviation of altitude | 18.0 | 8.5 | 10.2 | 20.5 | 8.7 | 11.7 | 12.9 | 8.7 |

**Table S4:** Yearly overview of the mean territory size (in ha) for great tits for the different subpopulations used in this study. Territory size is used as a measure of density, with higher mean territory sizes corresponding to lower density.

| **Year** | **b** | **c** | **cp** | **ex** | **mp** | **o** | **sw** | **w** |
| --- | --- | --- | --- | --- | --- | --- | --- | --- |
| 1980 | 2.07 | 1.00 | 3.80 | 3.42 | 1.74 | 2.61 | 3.06 | 2.75 |
| 1981 | 0.81 | 0.56 | 1.61 | 1.58 | 0.86 | 1.50 | 1.50 | 1.13 |
| 1982 | 1.43 | 0.89 | 1.92 | 2.32 | 2.45 | 2.20 | 2.12 | 1.79 |
| 1983 | 0.85 | 0.64 | 1.13 | 1.35 | 2.35 | 1.10 | 1.29 | 1.09 |
| 1984 | 1.11 | 1.29 | 2.64 | 2.20 | 4.79 | 1.87 | 2.16 | 1.02 |
| 1985 | 1.13 | 1.77 | 2.49 | 2.54 | 1.63 | 2.44 | 2.94 | 1.61 |
| 1986 | 0.88 | 0.82 | 2.09 | 2.49 | 1.04 | 2.31 | 2.55 | 1.40 |
| 1987 | 0.99 | 0.83 | 1.81 | 2.19 | 0.80 | 1.95 | 2.22 | 1.29 |
| 1988 | 0.75 | 0.56 | 1.60 | 1.70 | 0.68 | 1.82 | 2.31 | 1.15 |
| 1989 | 1.12 | 0.87 | 1.37 | 2.29 | 0.90 | 1.99 | 3.45 | 1.05 |
| 1990 | 1.30 | 0.75 | 1.38 | 2.18 | 4.84 | 3.68 | 3.66 | 1.38 |
| 1991 | 2.17 | 1.09 | 5.09 | 7.49 | 2.13 | 4.37 | 3.02 | 3.22 |
| 1992 | 1.62 | 1.16 | 1.72 | 2.42 | 3.79 | 1.97 | 2.83 | 1.17 |
| 1993 | 0.76 | 0.69 | 1.19 | 1.92 | 0.59 | 1.78 | 1.40 | 0.86 |
| 1994 | 0.8 | 0.69 | 1.29 | 1.87 | 0.72 | 1.24 | 1.41 | 0.75 |
| 1995 | 0.78 | 0.68 | 1.09 | 1.46 | 0.66 | 0.97 | 1.21 | 0.72 |
| 1996 | 0.67 | 0.49 | 0.96 | 1.22 | 0.59 | 1.08 | 1.39 | 0.65 |
| 1997 | 0.81 | 0.63 | 1.12 | 1.57 | 0.59 | 1.20 | 1.66 | 0.77 |
| 1998 | 0.9 | 0.64 | 1.12 | 1.40 | 0.55 | 1.53 | 1.46 | 0.78 |
| 1999 | 0.91 | 0.71 | 1.80 | 2.04 | 0.57 | 1.89 | 1.87 | 1.06 |
| 2000 | 0.86 | 0.49 | 1.82 | 2.50 | 0.63 | 2.23 | 2.01 | 1.12 |
| 2001 | 0.63 | 0.42 | 1.56 | 1.99 | 0.68 | 1.96 | 1.53 | 1.16 |
| 2002 | 0.61 | 0.42 | 1.13 | 2.05 | 0.51 | 1.6 | 1.21 | 0.91 |
| 2003 | 0.5 | 0.43 | 0.96 | 1.81 | 0.38 | 1.43 | 1.01 | 0.83 |
| 2004 | 0.76 | 0.47 | 1.75 | 1.85 | 0.57 | 1.58 | 1.63 | 0.90 |
| 2005 | 0.46 | 0.33 | 0.87 | 1.31 | 0.44 | 1.05 | 1.00 | 0.62 |
| 2006 | 0.69 | 0.42 | 0.97 | 1.32 | 0.53 | 1.1 | 1.18 | 0.62 |
| 2007 | 0.59 | 0.43 | 0.87 | 1.13 | 0.45 | 1.07 | 1.05 | 0.49 |
| 2008 | 0.56 | 0.48 | 0.97 | 1.23 | 0.51 | 1.16 | 0.93 | 0.58 |
| 2009 | 0.75 | 0.55 | 1.31 | 2.06 | 0.66 | 1.57 | 1.45 | 0.90 |
| 2010 | 0.64 | 0.48 | 1.16 | 1.72 | 0.55 | 1.41 | 1.23 | 0.70 |
| 2011 | 0.72 | 0.57 | 1.25 | 1.64 | 0.59 | 1.52 | 1.32 | 0.81 |
| 2012 | 0.62 | 0.41 | 1.03 | 1.46 | 0.49 | 1.26 | 1.03 | 0.78 |
| 2013 | 0.77 | 0.62 | 1.75 | 2.62 | 0.75 | 2.00 | 1.75 | 1.33 |
| 2014 | 0.67 | 0.59 | 1.58 | 2.28 | 0.60 | 1.86 | 1.01 | 1.06 |
| 2015 | 0.65 | 0.63 | 1.44 | 1.93 | 0.55 | 1.54 | 1.18 | 0.92 |
| 2016 | 1.02 | 1.14 | 1.55 | 1.86 | 0.60 | 1.83 | 1.68 | 0.96 |
| 2017 | 0.82 | 0.97 | 1.48 | 1.85 | 0.94 | 1.40 | 1.30 | 0.93 |
| 2018 | 1.33 | 0.84 | 1.37 | 1.95 | 0.92 | 1.34 | 1.65 | 1.10 |
| 2019 | 0.71 | 0.56 | 1.32 | 1.45 | 0.60 | 1.22 | 1.13 | 0.65 |
| 2020 | 0.92 | 0.74 | 1.94 | 3.14 | 0.71 | 1.80 | 1.81 | 0.97 |
| 2021 | 1.00 | 0.78 | 1.65 | 2.85 | 0.71 | 2.04 | 1.57 | 0.92 |
| 2022 | 0.99 | 0.62 | 1.69 | 2.57 | 0.99 | 2.06 | 1.83 | 1.10 |

**Table S5:** Yearly overview of the mean territory size (in ha) for blue tits for the different subpopulations used in this study. Territory size is used as a measure of density, with higher mean territory sizes corresponding to lower density.

| **year** | **b** | **c** | **cp** | **ex** | **mp** | **o** | **sw** | **w** |
| --- | --- | --- | --- | --- | --- | --- | --- | --- |
| 1980 | 0.73 | 0.64 | 3.76 | 3.06 | 0.55 | 1.4 | 1.46 | 1.19 |
| 1981 | 0.52 | 0.38 | 1.53 | 2.89 | 0.43 | 1.37 | 1.54 | 0.76 |
| 1982 | 0.66 | 0.56 | 1.53 | 3.01 | 2.93 | 1.64 | 2.34 | 0.96 |
| 1983 | 0.74 | 0.46 | 2.61 | 3.5 | 2.16 | 2.09 | 1.52 | 1.04 |
| 1984 | 1.03 | 0.42 | 2.05 | 3.28 | 1.26 | 1.31 | 1.74 | 1.26 |
| 1985 | 0.69 | 0.36 | 1.17 | 2.15 | 0.78 | 1.32 | 1.55 | 0.64 |
| 1986 | 0.89 | 0.4 | 2.01 | 2.98 | 0.88 | 1.92 | 2.73 | 0.95 |
| 1987 | 0.81 | 0.42 | 2.29 | 4.44 | 0.9 | 4.61 | 4.96 | 2.12 |
| 1988 | 0.55 | 0.39 | 3.48 | 4.33 | 0.56 | 2.45 | 2.19 | 0.74 |
| 1989 | 0.61 | 0.4 | 2.65 | 4.11 | 0.59 | 1.33 | 2.23 | 2.58 |
| 1990 | 0.49 | 0.32 | 2.42 | 1.78 | 0.7 | 1.09 | 1.32 | 0.79 |
| 2001 | 1.79 | 0.97 | 4.77 | 5.53 | 1.58 | 3.15 | 4.31 | 2.38 |
| 2002 | 3.07 | 2.19 | 6 | 5.22 | 2.65 | 3.95 | 4.26 | 2.14 |
| 2003 | 2.12 | 0.77 | 2.97 | 4.78 | 1.69 | 2.42 | 5.58 | 2.9 |
| 2004 | 0.57 | 0.65 | 1.22 | 4.33 | 0.81 | 1.72 | 2.55 | 1.47 |
| 2005 | 0.74 | 0.75 | 0.86 | 3.48 | 0.8 | 1.03 | 2.24 | 1.45 |
| 2006 | 0.7 | 4.12 | 8.62 | 10.85 | 0.76 | 2.24 | 2.41 | 8.56 |
| 2007 | 0.56 | 0.88 | 1.66 | 4.53 | 0.81 | 1.34 | 1.49 | 3.32 |
| 2008 | 0.58 | 0.44 | 1.36 | 5.25 | 0.67 | 10.02 | 2.75 | 2.35 |
| 2009 | 0.63 | 0.55 | 1.72 | 2.72 | 0.58 | 2.19 | 2.39 | 1.24 |
| 2010 | 0.42 | 0.36 | 0.62 | 1.48 | 0.65 | 0.82 | 1.31 | 0.57 |
| 2011 | 0.33 | 0.29 | 0.97 | 1.52 | 0.46 | 0.75 | 1.11 | 0.61 |
| 2012 | 0.45 | 0.42 | 0.7 | 1.54 | 0.51 | 0.8 | 1.53 | 0.51 |
| 2013 | 0.75 | 0.86 | 0.93 | 1.84 | 0.71 | 1.17 | 3.04 | 0.62 |
| 2014 | 0.68 | 0.45 | 0.75 | 1.5 | 0.67 | 0.88 | 1.77 | 0.55 |
| 2015 | 0.85 | 0.68 | 2.58 | 2.47 | 0.94 | 1.06 | 2.73 | 0.89 |
| 2016 | 0.79 | 0.43 | 0.68 | 1.67 | 0.94 | 0.81 | 1.27 | 0.6 |
| 2017 | 1.35 | 0.57 | 0.9 | 1.89 | 0.75 | 1.02 | 2.03 | 0.56 |
| 2018 | 0.89 | 0.44 | 0.85 | 1.77 | 0.73 | 0.94 | 2.02 | 0.57 |
| 2019 | 0.67 | 0.37 | 0.58 | 1.53 | 0.59 | 0.83 | 1.36 | 0.56 |
| 2020 | 0.4 | 0.3 | 0.55 | 1.16 | 0.4 | 0.69 | 0.97 | 0.4 |
| 2021 | 0.48 | 0.27 | 0.54 | 1.14 | 0.45 | 0.66 | 1.33 | 0.46 |
| 2022 | 0.51 | 0.35 | 0.58 | 1.18 | 0.81 | 0.68 | 1.14 | 0.42 |
| 2023 | 1.05 | 0.38 | 0.71 | 1.4 | 0.74 | 0.72 | 1.16 | 0.47 |

**Table S6:** The mean laying date and clutch size of the population for great tit and blue tit. Represented are also the standard deviations (σ) of the random effects and the h parameter, representing the proportion of the total unexplained variance that is explained by the different subpopulations.

| **Species** | **Great tit** | | **Blue tit** | |
| --- | --- | --- | --- | --- |
| **Phenotypic trait** | **Laying date** | **Clutch size** | **Laying date** | **Clutch size** |
| **Fixed effects (*β*)** | ***β*** | ***β*** | ***β*** | ***β*** |
| Intercept | **23.58 (21.42, 25.73)** | **8.47 (8.18, 8.76)** | **22.48 (20.00, 24.95)** | 9.88 (9.56, 10.20) |
| **Random effects (σ)** | **σ** | **σ** | **σ** | **σ** |
| Year | 6.32 (5.15, 7.85) | 0.70 (0.57, 0.87) | 6.55 (5.18, 8.38) | 0.65 (0.51, 0.84) |
| Subpopulation | 1.48 (0.93, 2.68) | 0.28 (0.17, 0.50) | 1.53 (0.96, 2.80) | 0.31 (0.19, 0.55) |
| Residual variance | 6.89 (6.77, 6.98) | 1.57 (1.55, 1.59) | 6.38 (6.29, 6.48) | 1.86 (1.83, 1.88) |
| **Quantified parameter** | ***h*** | ***h*** | ***h*** | ***h*** |
| Proportion of total variance | 0.02 | 0.02 | 0.03 | 0.02 |

**Supplementary materials: Dispersal**

While fledgling numbers are locality-based (recorded in a nest-box), recruitment numbers are more challenging, as the birds are free to move after they fledgling from the nest-box. If birds recruit outside of the population, this will reduce the recruit numbers. Moreover, if dispersal-rates are dependent upon the traits we estimate selection for, this could bias our selection estimates.

We had access to a limited dataset of both great tits (n=244 recoveries) and blue tits (n=196 recoveries) recorded outside of Wytham. A smaller proportion of these were ringed in nest-boxes, and could therefore be connected to broods with clutch size and laying date data (great tit n=87 from 86 different clutches; blue tit n=84 from 83 different clutches). The recoveries are spread in many directions and circumstances varied from re-captures at other places and newly dead birds killed by various sources (window-craches, cats etc.). The duration between recordings varied from 6 days up til almost 7 years (mean= 8 months and 11 days). To avoid restricting the data even further, we assumed that all birds found outside of Wytham would have recruited, irrespective of the circumstance of recovery. This is, of course, a simplifying assumption. However, it can still be informative for what birds tend to seek out of the population. Nevertheless, these results should be seen as complementary rather than conclusive on the role of dispersal.

**Table S7:** The effect of clutch size, laying date and their quadratic terms in addition to their interaction on the number of dispersers produced per clutch for great tits and blue tits. Both clutch size and laying date is standardized within the subpopulation-year. The model is based on a frequentist modelling approach (using the “lme4”-package). The 95% CI is marked in parentheses and statistically significant results are marked in bold.

| **Species** | **Great tit** | **Blue tit** |
| --- | --- | --- |
| **Fixed effects (β)** | ***β*** | ***β*** |
| Intercept | **-5.04 (-5.46, -4.61)** | **-5.50 (–6.16, –4.88)** |
| Clutch size | **0.34 (0.00, 0.69)** | 0.20 (–0.04, 0.45) |
| Quadratic clutch size | **-0.35 (–0.67, –0.03)** | 0.07 (–0.05, 0.19) |
| Laying date | 0.13 (–0.28, 0.55) | 0.20 (–0.20, 0.63) |
| Quadratic laying date | -0.31 (–0.76, 0.14) | 0.04 (–0.20, 0.29) |
| Clutch size: laying date | -0.47 (–1.09, 0.23) | 0.21 (–0.04, 0.50) |
| **Random effects (σ)** | **σ** | **σ** |
| Subpopulation: Intercept | 0.003 | 0.002 |
| Year: Intercept | 0.940 | 1.568 |
| Subpopulation-year: Intercept | 0.001 | 0.016 |
| **Sample size (n)** | **n** | **n** |
| Number of subpopulations | 8 | 8 |
| Number of years | 44 | 34 |
| Number of subpopulation-years | 352 | 271 |
| Number of observations | 10742 | 9338 |

**Table S8:** The effect of laying date, the quadratic component of laying date, clutch size, the quadratic component of clutch size, and the correlational selection of clutch size and laying date when modelling the number of fledgelings and number of recruits, respectively, for both great tit and blue tit. Here, also female identity is included as a random effect. The model is based on a frequentist modelling approach (using the “lme4”-package). The values are the model estimates followed by the 95% confidence intervals (CI). Followed by the estimates for the random effects, including the random slopes for clutch size and laying date within each random effect.

| **Species** | **Great tit** | | **Blue tit** | |
| --- | --- | --- | --- | --- |
| **Fitness trait** | **Fledglings** | **Recruits** | **Fledglings** | **Recruits** |
| **Fixed effects (*β*)** | ***β*** | ***β*** | ***β*** | ***β*** |
| Intercept | **1.86 (1.78, 1.95)** | **-0.59 (-0.83, -0.38)** | **1.96 (1.86, 2.06)** | **-1.43 (-1.81, -1.05)** |
| Clutch size | **0.19 (0.17, 0.21)** | 0.03 (-0.03, 0.09) | **0.16 (0.13, 0.18)** | -0.05 (-0.13, 0.02) |
| Quadratic clutch size | **-0.05 (-0.06, -0.04)** | **-0.09 (-0.12, -0.06)** | **-0.05 (-0.06, -0.04)** | **-0.08 (-0.13, -0.04)** |
| Laying date | -0.03 (-0.08, 0.02) | **-0.39 (-0.51, -0.29)** | **-0.13 (-0.20, -0.06)** | **-0.73 (-0.94, -0.50)** |
| Quadratic laying date | **-0.05 (-0.08, -0.05)** | **-0.19 (-0.24, -0.13)** | **-0.10 (-0.12, -0.07)** | **-0.31 (-0.43, -0.20)** |
| Laying date: clutch size | **-0.03 (-0.06, -0.01)** | **-0.09 (-0.17, -0.00)** | **-0.06 (-0.09, -0.02)** | **-0.15 (-0.28, -0.03)** |
| **Random effects (σ)** | **σ** | **σ** | **σ** | **σ** |
| Subpopulation: Intercept | 0.078 | 0.124 | 0.075 | 0.221 |
| Subpopulation: clutch size | 0.018 | 0.039 | 0.009 | 0.012 |
| Subpopulation: laying date | 0.031 | 0.009 | 0.050 | 0.115 |
| Subpopulation: clutch size: laying date | 0.017 | 0.024 | 0.004 | 0.033 |
| Year: Intercept | 0.204 | 0.705 | 0.244 | 1.000 |
| Year: clutch size | 0.021 | 0.125 | 0.056 | 0.121 |
| Year: laying date | 0.134 | 0.301 | 0.164 | 0.478 |
| Year: clutch size: laying date | 0.018 | 0.101 | 0.019 | 0.035 |
| Subpopulation-year: Intercept | 0.128 | 0.241 | 0.134 | 0.323 |
| Subpopulation-year: clutch size | 0.020 | 0.034 | 0.050 | 0.095 |
| Subpopulation-year: laying date | 0.099 | 0.267 | 0.131 | 0.263 |
| Subpopulation-year: clutch size: laying date | 0.065 | 0.192 | 0.094 | 0.212 |
| Female identity | 0.056 | 0.336 | 0.029 | 0.524 |
| **Sample size (n)** | **n** | **n** | **n** | **n** |
| Number of subpopulations | 8 | 8 | 8 | 8 |
| Number of years | 44 | 44 | 34 | 34 |
| Number of subpopulation-years | 352 | 352 | 268 | 268 |
| Number of observations | 9305 | 9243 | 7102 | 7102 |
| Number of females | 6167 | 6145 | 5509 | 5509 |

**Table S9:** The effect of edge-status on territory size of great tits in this population. Included is the model estimate and the 95% CI in parentheses. Statistically significant variables are marked in bold.

| **Fixed effects (β)** | **Territory size** |
| --- | --- |
| Intercept | **1.01 [0.86, 1.16]** |
| Edge-territory | **0.65 [0.62, 0.68]** |
| **Random effects (σ)** |  |
| Nestbox | 0.74 [0.71, 0.78] |
| Year | 0.47 [0.38, 0.58] |
| Residual variance | 0.67 [0.66, 0.68] |

**Table S10:** The effect of edge-status on the number of oak trees in each territory of great tits in this population. The model is based on a frequentist modelling approach (using the “lme4”-package). Included is the model estimate and the 95% CI in parentheses. Statistically significant variables are marked in bold.

| **Fixed effects (β)** | **Number of oaks in territory** |
| --- | --- |
| Intercept | **2.37 [2.30, 2.44]** |
| Edge-territory | -0.001 [-0.10, 0.10] |
| **Random effects (σ)** |  |
| Nestbox | 1.21 |

**Table S11:** The effect of laying date, the quadratic component of laying date, clutch size, the quadratic component of clutch size, and the correlational selection of clutch size and laying date when modelling the number of fledgelings and number of recruits, respectively, for both great tit and blue tit. This model is based on a frequentist modelling approach (using the “lme4”-package). The values are the model estimates followed by the 95% confidence intervals (CI). Followed by the estimates for the random effects, including the random slopes for clutch size and laying date within each random effect.

| **Species** | **Great tit** | | **Blue tit** | |
| --- | --- | --- | --- | --- |
| **Fitness trait** | **Fledglings** | **Recruits** | **Fledglings** | **Recruits** |
| **Fixed effects (*β*)** | ***β*** | ***β*** | ***β*** | ***β*** |
| Intercept | **1.79 (1.69, 1.89)** | **-0.64 (-0.88, -0.41)** | **1.90 (1.80, 2.01)** | **-1.34 (-1.72, -0.95)** |
| Clutch size | **0.19 (0.17, 0.21)** | 0.01 (-0.05,  0.07) | **0.16 (0.13, 0.18)** | -0.03 (-0.11, 0.05**)** |
| Quadratic clutch size | **-0.05 (-0.06, -0.05)** | **-0.10 (-0.13, -0.08)** | **-0.05 (-0.06, -0.04)** | **-0.12 (-0.16, -0.07)** |
| Laying date | -0.02 (-0.07, 0.04) | **-0.40 (-0.52, -0.29)** | **-0.11 (-0.19, -0.04)** | **-0.65 (-0.86, -0.44)** |
| Quadratic laying date | **-0.07 (-0.09, -0.05)** | **-0.18 (-0.23, -0.14)** | **-0.10 (-0.12, -0.07)** | **-0.32 (-0.40, -0.23)** |
| Laying date: clutch size | -0.02 (-0.06, 0.01) | **-0.10 (-0.19, -0.02)** | **-0.06 (-0.09, -0.03)** | **-0.18 (-0.31, -0.06)** |
| **Random effects (σ)** | **σ** | **σ** | **σ** | **σ** |
| Subpopulation: Intercept | 0.092 | 0.140 | 0.073 | 0.169 |
| Subpopulation: clutch size | 0.017 | 0.047 | 0.007 | 0.041 |
| Subpopulation: laying date | 0.029 | 0.014 | 0.041 | 0.087 |
| Subpopulation: clutch size: laying date | 0.029 | 0.039 | 0.000 | 0.058 |
| Year: Intercept | 0.240 | 0.709 | 0.270 | 1.051 |
| Year: clutch size | 0.020 | 0.114 | 0.058 | 0.139 |
| Year: laying date | 0.149 | 0.304 | 0.187 | 0.514 |
| Year: clutch size: laying date | 0.008 | 0.084 | 0.024 | 0.078 |
| Subpopulation-year: Intercept | 0.124 | 0.269 | 0.132 | 0.333 |
| Subpopulation-year: clutch size | 0.012 | 0.100 | 0.038 | 0.168 |
| Subpopulation-year: laying date | 0.087 | 0.303 | 0.099 | 0.340 |
| Subpopulation-year: clutch size: laying date | 0.030 | 0.171 | 0.059 | 0.277 |
| **Sample size (n)** | **n** | **n** | **n** | **n** |
| Number of subpopulations | 8 | 8 | 8 | 8 |
| Number of years | 44 | 44 | 34 | 34 |
| Number of subpopulation-years | 352 | 352 | 271 | 271 |
| Number of observations | 10477 | 10382 | 8986 | 8986 |

**Table S12:** The proportion of variation in selection for the different traits (CS= clutch size, LD = laying date and LD:CS=interaction between laying date and clutch size) explained by spatio-temporal, temporal and spatial components for the two models treating fledgling number and recruit number as fitness for each species. Here, the phenotypes are standardized to the population mean for the given year to reduce the impact of varying phenotypic distributions among the subpopulations, The parameters are based on a Bayesian model using the “Stan”-package. Presented here are the posterior medians and the 95% credible intervals. While the values for each trait per fitness trait and species should sum to 1, these results are based on a Bayesian approach and may therefore deviate slightly from this sum.

| **Species** | **Fitness trait** | **Source** | **LD** | **CS** | **LD:CS** |
| --- | --- | --- | --- | --- | --- |
| **Great tit** | **Fledglings** | **Spatio-temporal** | 0.45 (0.31, 0.64) | 0.83 (0.50, 0.97) | 0.84 (0.45, 0.97) |
|  |  | **Temporal** | 0.51 (0.35, 0.64) | 0.05 (0.00, 0,17) | 0.01 (0.00, 0.07) |
|  |  | **Spatial** | 0.03 (0.01,0.11) | 0.12 (0.02, 0.39) | 0.15 (0.03, 0.52) |
|  | **Recruits** | **Spatio-temporal** | 0.63 (0.54, 0.71) | 0.46 (0.25, 0.60) | 0.71 (0.43, 0.85) |
|  |  | **Temporal** | 0.37 (0.29, 0.46) | 0.18 (0.07, 0.32) | 0.20 (0.05, 0.46) |
|  |  | **Spatial** | 0.00 (0.00, 0.03) | 0.35 (0.25, 0.58) | 0.10 (0.01, 0.28) |
| **Blue tit** | **Fledglings** | **Spatio-temporal** | 0.45 (0.34, 0.61) | 0.66 (0.39, 0.86) | 0.97 (0.82, 1.00) |
|  |  | **Temporal** | 0.50 (0.36, 0.64) | 0.30 (0.11, 0.59) | 0.02 (0.00, 0.13) |
|  |  | **Spatial** | 0.03 (0.00, 0.16) | 0.02 (0.00, 0.22) | 0.00 (0.00, 0.08) |
|  | **Recruits** | **Spatio-temporal** | 0.30 (0.11, 0.54) | 0.76 (0.19, 0.99) | 0.83 (0.30, 0.99) |
|  |  | **Temporal** | 0.66 (0.40, 0.86) | 0.16 (0.01, 0.54) | 0.04 (0.00, 0.32) |
|  |  | **Spatial** | 0.03 (0.00, 0.21) | 0.06 (0.00, 0.60) | 0.11 (0.00, 0.63) |

**Table S13:** Estimated standard deviations (σ) of mean phenotypes (z̄) of clutch size and laying date, calculated both across all subpopulations and for the entire population for both great tits and blue tits. Included are also the standard deviations of the estimated optimums (θ) for fledgling- and recruit production, respectively.

| Species | Variable | Clutch size | | Laying date | |
| --- | --- | --- | --- | --- | --- |
|  |  | Subpopulation | Population | Subpopulation | Population |
| Great tit | *σ_z̄_* | 0.85 | 0.71 | 6.76 | 6.38 |
|  | *σ_θ_* (fledglings) | 0.46 | 0.33 | 9.87 | 8.97 |
|  | *σ_θ_* (recruits) | 1.05 | 0.85 | 8.31 | 6.56 |
| Blue tit | *σ_z̄_* | 0.89 | 0.63 | 6.87 | 6.52 |
|  | *σ_θ_* (fledglings) | 1.20 | 0.98 | 8.28 | 7.96 |
|  | *σ_θ_* (recruits) | 1.25 | 0.97 | 6.71 | 6.21 |

**Table S14:** The effect of clutch size, laying date and their quadratic terms in addition to the effect of breeding in an edge-territory on the number of dispersers produced for great tits and blue tits. Clutch size and laying date are standardized within the edge-status-year. The model is based on a frequentist modelling approach (using the “lme4”-package). The 95% CI is marked in parentheses and statistically significant results are marked in bold.

| **Species** | **Great tit** | **Blue tit** |
| --- | --- | --- |
| **Fixed effects (β)** | ***β*** | ***β*** |
| Intercept | **–5.08 (–5.54, –4.62)** | **–5.76 (–6.46, –5.11)** |
| Clutch size | **0.29 (0.01, 0.59)** | 0.24 (–0.00, 0.49) |
| Quadratic clutch size | –0.15 (–0.37, 0.08) | –0.02 (–0.13, 0.08) |
| Laying date | –0.09 (–0.43, 0.25) | 0.11 (–0.32, 0.50) |
| Quadratic laying date | –0.01 (–0.24, 0.22) | -0.01 (–0.30, 0.30) |
| Edge-territory | –0.17 (–0.64, 0.27) | **0.62 (0.19, 1.10)** |
| **Random effects (σ)** | **σ** | **σ** |
| Edge-status-year: Intercept | 0.000 | 0.000 |
| Year: Intercept | 0.925 | 1.536 |
| **Sample size (n)** | **n** | **n** |
| Number of years | 43 | 34 |
| Number of edge-status-years | 86 | 68 |
| Number of observations | 10481 | 9388 |

**Table S15:** Selection estimates in great tit and blue tit for clutch size, the quadratic component of clutch size, laying date and the quadratic component of laying date for the models of number of fledglings and number of recruits, respectively. Further is the effect of edge-territories on the fitness variable and the interaction terms with edge-territories and the predictors and their quadratic components. Here, only breeding events with known mother identity are included and used as a random effect. The model is based on a frequentist modelling approach (using the “lme4”-package).  The values are the model estimates followed by the 95 % confidence intervals (CI). Further are the random effects including random slope between the random effects and the predictors.

| **Species** | **Great tit** | | **Blue tit** | |
| --- | --- | --- | --- | --- |
| **Fitness parameter** | **Fledglings** | **Recruits** | **Fledglings** | **Recruits** |
| **Fixed effects (*β*)** | ***β*** | ***β*** | ***β*** | ***β*** |
| Intercept | **1.87 (1.70, 1.94)** | **-0.80 (-1.01, -0.61)** | **1.93 (1.83, 2.03)** | **-1.24 (-1.61, -0.82)** |
| Clutch size | **0.22 (0.20, 0.23)** | **0.10 (0.04, 0.16)** | **0.19 (0.16, 0.22)** | 0.04 (-0.03,  0.11) |
| Quadratic clutch size | **-0.05 (-0.06, -0.04)** | **-0.07 (-0.10, -0.04)** | **-0.05 (-0.06, -0.03)** | -0.04 (-0.09,  0.01) |
| Laying date | -0.03 (-0.07, 0.02) | **-0.42 (-0.54, -0.30)** | **-0.10 (-0.18, -0.03)** | **-0.74 (-0.97, -0.53)** |
| Quadratic laying date | **-0.05 (-0.06, -0.03)** | **-0.12 (-0.19, -0.06)** | **-0.07 (-0.09, -0.05)** | **-0.23 (-0.34, -0.11)** |
| Edge-territories | -0.02 (-0.04, 0.01) | **-0.13 (-0.21, -0.05)** | **-0.06 (-0.09, -0.02)** | **-0.16 (-0.29, -0.03)** |
| Altitude | -0.00 (-0.00, 0.00) | **0.00 (0.00, 0.00)** | 0.00 (-0.00, 0.00) | -0.00 (-0.00,  0.00) |
| Territory size | **0.03 (0.02, 0.04)** | **0.09 (0.06, 0.12)** | 0.00 (-0.01, 0.01) | 0.03 (-0.02,  0.07) |
| Clutch size: Edge-territories | **-0.03 (-0.05, -0.00)** | -0.06 (-0.14, 0.01) | -0.03 (-0.07, 0.00) | -0.05 (-0.17,  0.08) |
| Quadratic clutch size: Edge-territories | 0.01 (-0.01, 0.02) | -0.00 (-0.05, 0.05) | 0.01 (-0.01, 0.03) | 0.01 (-0.07,  0.09) |
| Laying date: Edge-territories | -0.02 (-0.05, 0.01) | -0.01 (-0.12, 0.10) | -0.07 (-0.13, -0.00) | -0.02 (-0.33,  0.26) |
| Quadratic laying date: Edge-territories | 0.00 (-0.01, 0.02) | 0.03 (-0.05, 0.12) | 0.01 (-0.03, 0.04) | -0.04 (-0.22,  0.15) |
| **Random effects (σ)** | **σ** | **σ** | **σ** | **σ** |
| Edge-statusyear: Intercept | 0.040 | 0.084 | 0.036 | 0.000 |
| Edge-statusyear: clutch size | 0.018 | 0.071 | 0.032 | 0.048 |
| Edge-statusyear: laying date | 0.015 | 0.087 | 0.092 | 0.395 |
| Year: Intercept | 0.201 | 0.507 | 0.239 | 0.965 |
| Year: clutch size | 0.005 | 0.083 | 0.056 | 0.014 |
| Year: laying date | 0.129 | 0.313 | 0.183 | 0.135 |
| Female identity | 0.100 | 0.370 | 0.243 | 0.545 |
| **Sample size (n)** | **n** | **n** | **n** | **n** |
| Number of years | 43 | 43 | 34 | 34 |
| Number of edge-status-years | 86 | 86 | 68 | 68 |
| Number of females | 6017 | 5995 | 4578 | 4078 |
| Number of observations | 9040 | 8979 | 5783 | 5783 |
